# Climate and hybridization shape stomatal trait evolution in *Populus*

**DOI:** 10.1101/2025.07.21.665994

**Authors:** Michelle Zavala-Paez, Stephen Keller, Jason Holliday, Matthew C. Fitzpatrick, Jill Hamilton

## Abstract

- Stomata play a critical role in regulating plant responses to climate. Where sister species differ in stomatal traits, interspecific gene flow can influence the evolutionary trajectory of trait variation, with consequences to adaptation.
- Leveraging six latitudinally-distributed transects spanning the natural hybrid zone between *Populus trichocarpa–P. balsamifera*, we used whole genome resequencing and replicate common garden experiments to test the role that interspecific gene flow and selection play to stomatal trait evolution.
- While species-specific differences in the distribution of stomata persist between *P. balsamifera* and *P. trichocarpa*, hybrids on average resembled *P. trichocarpa*. Admixture mapping identified several candidate genes associated with stomatal trait variation in hybrids including *TWIST*, a homolog of *SPEECHLESS* in *Arabidopsis*, that initiates stomatal development via asymmetric cell divisions. Geographic clines revealed candidate genes deviating from genome-wide average patterns of introgression, suggesting restricted gene flow and the maintenance of adaptive differences. Climate associations, particularly with precipitation, indicated selection shapes local ancestry at candidate genes across transects.
- These results highlight the role of climate in shaping stomatal trait evolution in *Populus* and demonstrate how interspecific gene flow creates novel genetic combinations that may enhance adaptive potential in changing environments.

## Introduction

Hybridization is an important source of variation that enhances adaptive evolution by creating new recombinant genotypes and increasing standing genetic variation (Rieseberg & Wendel, 1993; Burke & Arnold, 2001; Brauer *et al*., 2023). The movement of alleles from one species into the genetic background of another via repeated backcrossing can facilitate adaptive evolution (Hamilton & Aitken, 2013; Hamilton *et al*., 2013; Hamilton & Miller, 2016; Suarez-Gonzalez *et al*., 2018b,c; Menon *et al*., 2021). This is particularly notable in plant species, where adaptive introgression has been associated with traits related to water use that are critical to maintaining physiological function under changing environmental conditions (Welch & Rieseberg, 2002; Campbell, 2004; Wu & Campbell, 2007; Whitney *et al*., 2010; Campbell & Wendlandt, 2013; Suarez-Gonzalez *et al*., 2018c). Despite potential benefits from introgression, fine-scale assessments of the genetic variation underlying adaptive trait variation and the role of extrinsic selection to introgression across different genomic backgrounds remains limited. This study aims to understand how natural selection shapes the movement of genetic variation across species boundaries that is essential for predicting how recombinant genotypes may respond to climate change.

Stomatal traits are fundamental to photosynthesis, transpiration, and overall water balance, and exhibit substantial variation across species (Hetherington & Woodward, 2003; Drake *et al*., 2013; Marek *et al*., 2022; Chen *et al*., 2024). These microscopic pores on the leaf surface regulate CO₂ uptake and water vapor loss, making them key targets of selection under changing environmental conditions (Chen *et al*., 2024). Stomata have evolved variation in size, density, and distribution across upper and lower leaf surfaces to optimize trade-offs associated with carbon uptake, water-use efficiency, and pathogen exposure across environments (Hetherington & Woodward, 2003; Dittberner *et al*., 2018; Chen *et al*., 2024). Natural hybrid zones provide ideal systems to understand how varying genomic and environmental backgrounds shape the broad and fine-scale evolutionary trajectory of stomatal traits. Genomic recombination via hybridization and subsequent introgression can produce new heritable phenotypes, including traits intermediate or in excess of the range observed in either parent species that may be beneficial for adaptation under changing environmental conditions (Campbell, 2004; Campbell & Wendlandt, 2013; Hamilton & Miller, 2016; Janes & Hamilton, 2017). Thus, identifying genes underlying stomatal variation and tracking their movement across species boundaries may reveal how interspecific gene flow facilitates adaptive trait evolution required for climate adaptation.

One approach to assessing the impact of gene flow into a contact zone and selection against recombination in a hybrid zone is geographic cline analysis, which investigates spatial gradients in traits or allele frequencies where species interbreed (Barton, 1979; Barton & Gale, 1993). Comparing geographic clines across repeated hybrid zones for traits, genome-wide ancestry, and candidate genes underlying adaptive traits allows an evaluation of how gene flow between species varies across genomic and environmental backgrounds (Stankowski *et al*., 2017; Schield *et al*., 2024). However, as clines may arise from either intrinsic or extrinsic selection, geographic cline analysis alone cannot distinguish the mechanisms facilitating or limiting genetic exchange (Hamilton & Aitken, 2013; Janes & Hamilton, 2017; Capblancq & Després, 2020). To address this gap, environmental associations may be used to identify the role extrinsic forces may play in adaptive introgression. Integrating geographic cline analyses with environmental associations can help identify traits or genes that move across both environmental and genomic backgrounds and are critical to adaptation under future climates.

*Populus* is an evolutionary and ecological model tree that has become an invaluable system for understanding how interspecific gene flow and natural selection shape trait variation (Taylor, 2002; Lexer *et al*., 2004; Buerkle & Lexer, 2008). In this study, we focus on the natural hybrid zone between *P. trichocarpa* × *P. balsamifera*, two sister species that have evolved in response to contrasting environmental conditions. The geographic distribution of *P. trichocarpa* comprises mild, moist environments of the Pacific Northwest, in contrast to *P. balsamifera,* whose distribution is confined to colder, drier boreal regions that experience greater seasonal extremes (Suarez-Gonzalez *et al*., 2018b,a). These environmental differences have contributed to the evolution of species-specific stomatal trait variation. On average, *P. trichocarpa* has large stomata on both the upper (adaxial) and lower (abaxial) leaf surfaces, a pattern consistent with amphistomaty (McKown *et al*., 2014, 2019). In contrast, *P. balsamifera* typically exhibits smaller, more densely packed stomata that are mostly restricted to the lower leaf surface, reflecting hypostomaty (Soolanayakanahally *et al*., 2009; Pointeau & Guy, 2014). Previous work has indicated that the movement of genomic regions from *P. balsamifera* into the genomic background of *P. trichocarpa* have contributed to adaptive trait evolution linked to persistence in colder and drier environments (Suarez-Gonzalez *et al*., 2018b). However, less is known about how genetic variation underlying stomatal traits moves across species boundaries (but see, Fetter & Keller, 2023), or how climate may influence that movement.

Here, we identify genetic variation underlying stomatal traits and compare the broad- and fine-scale role of geography and climate in shaping the movement of genetic variation across repeated contact zones between *P. trichocarpa* and *P. balsamifera*. Specifically we ask: 1) How do hybrids vary in stomatal traits relative to *P. trichocarpa* and *P. balsamifera*? 2) To what extent do stomatal traits vary across repeated contact zones, and are these patterns shaped by local climatic gradients? 3) What genes underlie stomatal trait variation, and how does the environment interact to influence the extent and direction of introgression for candidate genes across contact zones? This work identifies key candidate genes related to stomatal trait variation in *Populus* and their relationships to climate gradients providing a basis to predict whether and how genetic variation underlying adaptive traits may shift across species and environmental gradients under climate change.

## Material and Methods

### Study Approach

#### Sample collection and library preparation

In this study, we analyzed 574 genotypes collected within the natural contact zone between *Populus trichocarpa* × *P. balsamifera* hybrid zone (Fig. 1, Table S1; see Bolte *et al*., 2024 for full details). Dormant vegetative cuttings were sampled from six contact zones spanning a latitudinal gradient, referred to here as Alaska, Cassiar, Chilcotin, Jasper, Crowsnest, and Wyoming transects, respectively. Cuttings were propagated clonally under greenhouse conditions (24 °C day/15.5 °C night, no supplemental lighting) at the Virginia Tech Reynold’s Homestead Forest Resource Research Center (FRRC) in Stuart, VA, USA. Once established, 100 mg of fresh leaf tissue was collected from each genotype for DNA extraction and whole-genome resequencing. Methods for propagation, DNA extraction, library preparation, sequencing, and quality filtering are fully described in Bolte et al. 2024. Genomic libraries were sequenced on an Illumina NovaSeq 6000 using an S4 flow cell in 2 × 150 bp paired-end format, with 64 samples per lane. Illumina reads for each genotype were aligned to the *Populus trichocarpa* reference genome (v4.0), generating SAM files that were subsequently converted to BAM format using SAMtools (Li *et al*., 2009). Individual gVCF files were generated using the HaplotypeCaller algorithm in GATK v3.7 and subsequently combined into a single VCF file using the GenotypeGVCFs function. The initial dataset included ∼82 million variants and was filtered based on mapping quality (MQ < 40.00), strand bias (FS > 40.000, SOR > 3.0), mapping quality rank sum (MQRankSum < –12.500), read position bias (ReadPosRankSum < –8.000), and depth of coverage (QD < 2.0). INDELs and SNPs with more than two alternate alleles were excluded and then filtered to remove SNPs with a minor allele frequency below 5% and more than 10% missing data, resulting in a final dataset of 7,361,750 biallelic SNPs for downstream analyses. Additionally, thirty individuals were excluded based on ancestry estimates from Bolte et al. (2024) as their genomic composition did not correspond to either parental species.

**Figure 1.**
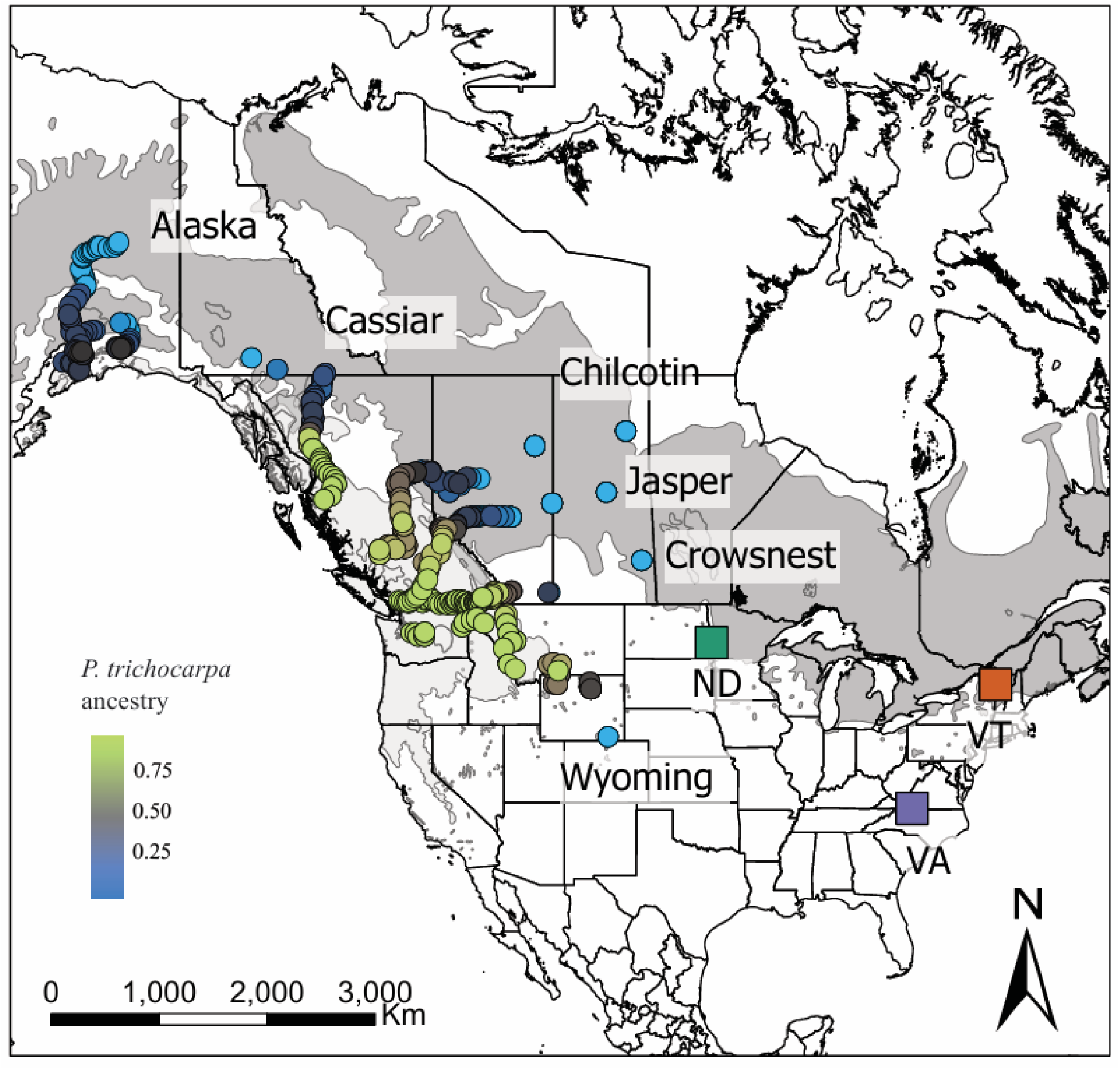
Genotypes were collected across six east-west contact zones within the natural hybrid zone between *P. trichocarpa* and *P. balsamifera*. Light and dark gray represent the distribution range of *P. trichocarpa* and P*. balsamifera*, respectively. Georeferenced sampling locations are color coded by genomic ancestry (Mead et al, 2025; K = 2). Clonally replicated genotypes were planted at the common gardens in Stuart, VA (purple square), Burlington, VT (orange square), and Fargo, ND (green square).

#### Common gardens design

In March 2020, the 574 clonally propagated rooted *Populus* cuttings were planted across three common garden environments (Fig. 1). The common gardens, located at the FRRC in Stuart, VA (36° 37’ N and 80° 09’ W, elevation 359 m), Burlington, VT (44° 26’ N and 73° 11’ W, elevation 130 m) and Fargo, ND (46° 53’ N and 96° 48’ W, elevation 272 m) were established spanning both latitudinal and longitudinal gradients in temperature and precipitation. At each site, genotypes were planted in a randomized complete block design with three replicates per genotype, resulting in a total of 1,722 individuals per site.

#### Quantifying stomatal traits in replicated common gardens

During the summer of 2022, stomatal traits and associated measures were assessed for each genotype across three common garden experiments (see Table1 for list). For each genotype, the first fully expanded leaf on the dominant shoot was used to assess stomatal conductance, size, density, and distribution on both the adaxial (upper) and abaxial (lower) leaf surfaces. This leaf was selected to minimize variation due to leaf age and environmental effects between samples (Fetter *et al*., 2021). Stomatal conductance (g_sw_, mmol m^−2^ s^−1^) was measured using a LI-600 porometer in three non-overlapping areas per each fully expanded leaf without detaching the leaf from the plant. Individual g_sw_ values represent the average of these three measurements.

Following g_sw_ measurements, leaves were collected to assess stomatal density, size, and distribution. A layer of Newskin liquid bandage was applied to the upper and lower leaf surfaces to make two stomatal impressions to capture variation across leaf surfaces. Stomatal impressions from one leaf per individual were mounted on slides without a cover slip, and images were captured using an Olympus BX-53 microscope equipped with an Olympus DP23 digital camera. Micrographs were standardized to a 0.37 × 0.25 mm grid for analysis. Stomatal size traits were measured and analyzed independently for each leaf surface using ImageJ (Abràmoff *et al*., 2004). These included adaxial (G_U_, µm) and abaxial (G_L_, µm) guard cell length, as well as adaxial (P_U_, µm) and abaxial (P_L_, µm) pore length. Stomatal measurements were obtained by overlaying four equally spaced lines across each micrograph, selecting five stomata per image. Individual values represent the average of five measurements per leaf surface. Stomatal density was estimated as the number of stomata per unit leaf area for the adaxial (*D_U_*, mm⁻²) and abaxial (*D_L_*, mm⁻²) leaf surfaces using automated counts in LeafNet (Li *et al*., 2022). To validate automated density measures, micrographs from 100 random genotypes were manually counted and measured. A strong correlation was observed between automated and manual counts (ρ = 0.95, *p* < 0.05, Fig. S1); therefore, automated values generated by LeafNet were used for all genotypes. Total stomatal density (D_T_) was also calculated as the sum of D_U_ and D_L_, representing the overall number of stomata per unit leaf area. To estimate the degree of amphistomy, stomatal ratio (SR) was calculated as the ratio of adaxial stomatal density divided by the total stomatal density. SR values range from 0 to 1, with 0 indicating stomata are present only on the abaxial surface, 0.5 indicating equal density on both surfaces, and 1 indicating stomata are exclusive to the adaxial surface (Fetter *et al*., 2021). To capture variation in adaxial stomatal occurrence (O_U_) among genotypes we assessed the presence (1) or absence (0) of adaxial stomata.

In addition to stomatal trait variation, intrinsic water-use efficiency (δ¹³C, ‰) was assessed using the second fully expanded leaf. Leaves were collected and dried at 60°C until a constant mass was reached. Dried leaf tissue was then homogenized into a fine powder using a TissueLyser II (Qiagen, Hilden, Germany). Approximately 2–3 mg of the homogenized tissue was placed into a tin capsule (Costech, Valencia, CA, USA) for analysis. Carbon isotope composition (δ¹³C, ‰) was measured at the Central Appalachians Stable Isotope Facility (CASIF), Appalachian Laboratory (Frostburg, Maryland, USA).

#### Comparing variation in stomatal traits among hybrid genotypes relative to parental species

To test whether hybrids exhibit intermediate, transgressive, or parental-like stomatal traits, we first estimated the genetic values of each genotype for each trait by calculating best linear unbiased predictors (BLUPs) using R 4.3.1 (R Core Team 2024). BLUPs were estimated for each trait for each genotype across the three common gardens experiments using the following linear mixed model.

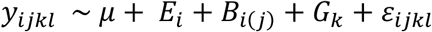

Where *y_*i*jkl_* is the observed trait value for clonal replicate *l*, *E*_*i*_ is the fixed effect of the common garden experiment, *B*_*i*(*i*)_ is the random effect of block *j* nested within garden *i*, *G*_*i*_ is the random effect of genotype, and *ɛ_ijkl_* is the residual term. We then extracted the genotype random effects (*G_k_*), which represents each genotype’s estimated genetic contribution to the trait after accounting for garden and block effects for use in downstream analyses.

Individuals were classified into three genotypic classes based on admixture proportions from Mead *et al*., (2025) to evaluate species-specific and hybrid differences in stomatal traits. Using ancestry estimates (*K* = 2), genotypes with Q > 0.98 were assigned to *P. trichocarpa* (*n* = 132), and those with Q < 0.02 to *P. balsamifera* (*n* = 91). Remaining genotypes (*n* = 323) were categorized as hybrids. To test for differences in stomatal traits among *P. trichocarpa*, *P. balsamifera*, and their hybrids, we first assessed homogeneity of variance using Levene’s test in R 4.3.1 (RStudio Team 2024). For traits that met the assumptions of homogeneity, we used one-way ANOVA followed by Tukey’s HSD for post hoc comparisons. For traits that violated this assumption, we applied the non-parametric Kruskal–Wallis test with Dunn’s post hoc comparisons. Four out of eleven stomatal traits, including g_sw_, D_U_, SR, and O_U_, did not exhibit homogeneity of variance across groups and were therefore analyzed using the non-parametric tests.

#### Admixture mapping to identify the genetic basis of stomatal variation

To identify the genetic basis of stomatal trait variation and water-use efficiency, we performed admixture mapping using local ancestry inferred with Loter (Dias-Alves *et al*., 2018) and phenotype-genotype association conducted with GEMMA (Zhou & Stephens, 2014). Local ancestry inference relies on phased haplotypes from reference parental genotypes to infer the ancestral origin of specific genomic regions in admixed genotypes (Dias-Alves *et al*., 2018) Prior to local ancestry inference, the VCF file was phased and missing data imputed using BEAGLE (Browning *et al*., 2018, 2021). Parental reference genotypes for local ancestry inference were the same as those used for trait comparisons as defined above. These phased reference haplotypes were used to infer local ancestry across the genomes of 323 admixed individuals.

To associate genomic variation with trait variation we used Univariate Linear Mixed Models implemented in GEMMA v.0.94.1 (Zhou & Stephens, 2014). A relatedness matrix (estimated with the -gk 1 option) was included as a covariate to account for relatedness among individuals. Global ancestry, estimated as the average local ancestry across all loci, was also included as a covariate to account for population structure (Fetter & Keller, 2023). To account for multiple testing, the significance threshold was adjusted using the admixture burden (0.05/2719), which estimates the number of independently recombining chromosomal segments within a genotype. Admixture burden was estimated following the method described by Shriner *et al*., (2011). Specifically, an autoregressive (AR) model was fitted to the local ancestry sequence of each genotype, and the spectral density of local ancestry values at frequency zero was estimated using the spectrum0.ar function from the coda package in R 4.3.1. The number of independent ancestry blocks per individual was then summed and averaged across all individuals to estimate the mean effective number of independent tests in our dataset (Shriner *et al*., 2011). This value was used to adjust the significance threshold for admixture mapping. Manhattan plots were generated using the qqman package in R to visualize admixture mapping results, with significance assessed based on the admixture burden threshold (Turner, 2018).

To identify candidate genes associated with stomatal traits, we mapped loci that passed the admixture burden threshold to the *P. trichocarpa* v4.1 gene annotation. To account for upstream regulatory regions, gene boundaries were extended by 2 kb upstream (positive strand) or downstream (negative strand). We performed manual BLAST searches on TAIR (www.arabidopsis.org) to identify orthologous genes in *Arabidopsis thaliana*, and selected the best hits based on sequence similarity. When multiple genes were located within the same window, all were initially retained as candidates. Each gene was then evaluated for its potential role in stomatal development and function. Candidate gene-specific ancestry values were extracted from the ancestry matrix using gene positions from the *P. trichocarpa* v4.1 reference genome, including a ±500 bp flanking region. For each admixed individual, locus-specific ancestry at each gene was classified as homozygous *P. trichocarpa* (1), homozygous *P. balsamifera* (0), or heterozygous (0.5).

#### Geographic clines to identify barriers to gene flow for stomatal traits, candidate genes and genome-wide ancestry

To quantify the extent and direction of introgression for stomatal traits and associated candidate genes, we fitted geographic clines separately for each of the six contact zones. Geographic distances from the coast for each genotype within each transect were calculated using the Haversine formula and scaled between 0 and 1 to allow comparisons across contact zones (see Supplementary Methods). Geographic cline parameters, including center and width (1/ maximum slope, Derryberry *et al*., 2014), were estimated using genotype-specific BLUP values for traits and ancestry genotypes for candidate genes using HZAR (Derryberry *et al*., 2014).

For each trait, six different cline models were fit for each contact zone using different tails (both, left, right, mirrored, none, null). Each model was run with a burn-in of 100,000 iterations followed by 500,000 Monte Carlo iterations to estimate posterior distributions of cline parameters. The best-supported model for each trait in each contact zone was identified using corrected Akaike Information Criterion (AICc), selecting the model with the lowest AICc score. Cline parameter estimates based on the best-fit model were compared across contact zones.

For each candidate gene, ten different cline models using different combinations of scaling (free or fixed) and tails (both, left, right, mirrored, or none), in addition to the null model (no cline) were fit for each geographic transect. The inclusion of scaling and tail combinations allowed us to capture the full range of potential variation in ancestry genotypes across the contact zones. As with the trait clines, each gene cline model was run with 100,000 burn-in iterations and 500,000 Monte Carlo iterations. AICc scores were used to select the best-fitting model for each candidate gene and contact zone, and cline parameter estimates from these models were compared across contact zones. We compared fine-scale introgression of individual candidate genes with genome-wide ancestry, therefore cline parameters were estimated for genome-wide ancestry within each contact zone.

#### Modeling the influence of climate on stomatal traits, candidate genes and genome-wide ancestry

To complement geographic cline analyses, which identifies barriers to gene flow across space, we modeled climatic clines to test whether climate predicts variation in stomatal traits, candidate genes, and genome-wide ancestry within each contact zone. Twenty-five annual climate normals (1961–1990) were obtained from ClimateNA (Wang *et al*., 2016) using the latitude, longitude, and elevation of each genotype origin. Due to correlations among climate variables, we conducted a principal component analysis (PCA) using the prcomp function in R 4.3.1 to reduce multicollinearity and summarize climatic variation across genotypes based on climate of origin. PC1 explained 55% of climate variation, and captured temperature-related gradients, with high loadings for mean annual temperature and degree-days above 5 °C and 18 °C. PC2 explained 26% of climate variation, and captured moisture-related gradients, with high loadings for mean annual precipitation, climate moisture index, and relative humidity (Table S1, Figure S2). Given that PC1 and PC2 encompassed the major axes of climate variation, subsequent analyses focused on these two PC axes.

For each trait and contact zone, linear regressions were fit to quantify the relationship between climate and trait variation. Genotype-specific BLUP values were used as the response variable, and the two first climate principal components (PC1 and PC2) as predictors within the lm function in R 4.3.1 (R Core Team, 2024) following the equation:

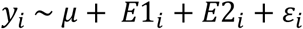

where *y*_*i*_ is the BLUP-estimated trait value for genotype *i*, μ is the intercept, and *E*1 and *E*2 represent the scores for PC1 (temperature-related gradient) and PC2 (moisture-related gradient), respectively, and *ɛ*_*i*_ is the residual error term. An interaction term between PC1 and PC2 was included in initial models, but was not significant for the majority of trait–transect combinations. To assess whether the direction and strength of climate–trait relationships varied across transects, slopes were compared across contact zones.

For each candidate gene, we tested whether variation in gene ancestry was associated with climatic gradients using logistic regression models implemented within the glm() function in R 4.3.1. We fit a single model for each gene in each transect, including both PC1 and PC2 as well as their interaction (PC1 × PC2) as predictors following the equation:

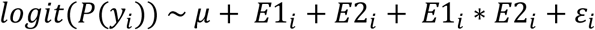

Where, where *y*_*i*_ is the BLUP-estimated trait value for genotype *i*, μ is the intercept, and *E*1 and *E*2 represent the scores for PC1 (temperature-related gradient) and PC2 (moisture-related gradient), respectively. *E*1_*i*_ ∗ *E*2_*i*_ represents the interaction term between PC1 and PC2, and *ɛ*_*i*_ is the residual error. To compare climatic associations between candidate genes and the genome-wide average, the same model was fit using genome-wide ancestry as the response variable. Slope values from candidate gene models were compared to those from the genome-wide models within each contact zone to evaluate differences in the strength and direction of climate–ancestry relationships. Slopes were also compared across contact zones for each candidate gene to evaluate whether climate–gene ancestry relationships vary.

## Results

### Species-specific and hybrid variation in stomatal traits between *Populus balsamifera* and *P. trichocarpa*

Parental genotypes of *Populus trichocarpa* and *P. balsamifera* differed significantly in several stomatal traits, with hybrids exhibiting intermediate values or resembling one of the parental species (Fig. 2A, Table 1). On average, *P. trichocarpa* exhibited reduced abaxial guard cell length (G_L_) compared to *P. balsamifera* (*p* < 0.05, Fig. 2A, Table S2), while adaxial guard cell length (G_U_) did not differ significantly between parental species (Fig. S3). In hybrids, G_L_ was similar to *P. balsamifera* (*p* > 0.05, Fig. 2A); however, hybrids exhibited a broader range in G_L_ than either parental species. Overall, G_U_ (F_2,496_ = 9.36, *p* > 0.05) and adaxial pore length (P_U_; F_2,386_ = 2.91, *p* > 0.05, Fig. S3) were not significantly different between parental and hybrid genotypes. In contrast, abaxial pore length (P_L_) differed significantly among genotype classes (*F*_2, 496_ = 4.39, *p* < 0.05, Fig. S3); post hoc comparisons showed that hybrid genotypes had significantly greater P_L_ than *P. trichocarpa* (*p* < 0.05) but did not differ from *P. balsamifera* (*p* > 0.05). Hybrids also exhibited a greater range of P_L_ values relative to parental species.

**Figure 2.**
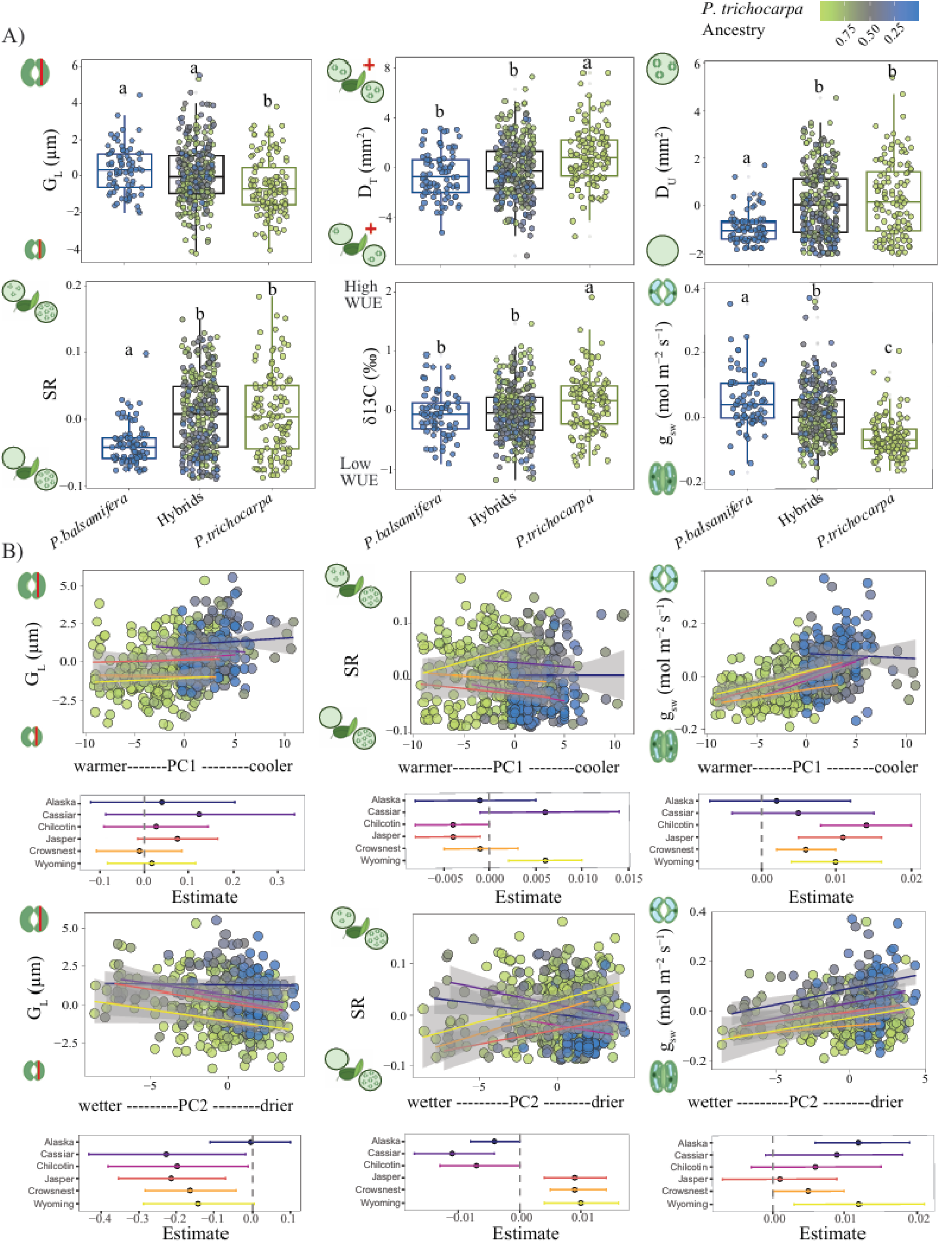
A) Boxplots showing best linear unbiased predictors (BLUPs) for abaxial guard cell length (G_L_), total stomatal density (D_T_), adaxial stomatal density (D_U_), stomatal ratio (SR), intrinsic water-use efficiency (δ¹³C), and stomatal conductance (g_sw_), for *P. balsamifera*, hybrid, and *P. trichocarpa* genotypes. Each point represents a genotype, and colors indicate the proportion of *P. trichocarpa* genomic ancestry. Letters denote statistically significant differences among groups based on post-hoc comparisons (*p* < 0.05). B) Relationship between genotype BLUPs values for abaxial guard cell length (G_L_), stomata ratio (SR), stomatal conductance (g_sw_), and the first two principal components of climate at genotypes’ origin. PC2 reflects a gradient from wetter to drier environments (increasing values = drier), while PC1 reflects a temperature gradient (increasing values = warmer). Shaded areas indicate 95% confidence intervals for the fitted linear regression. Lower panels show trait-transects slopes (±95% CI) extracted from linear models fit separately within each transect.

**Table 1.**
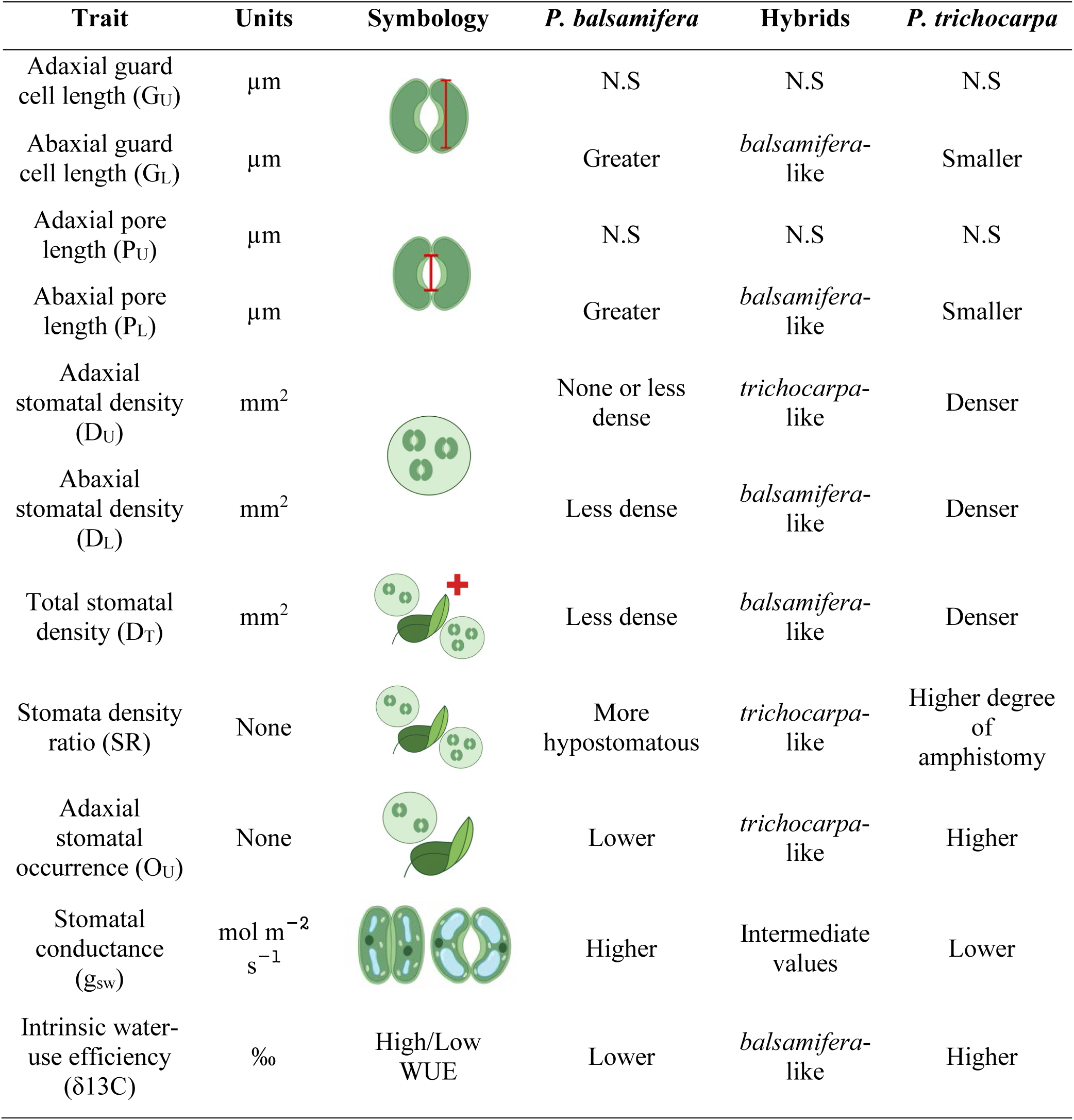
Summary of stomatal traits and associated measures assessed across the common gardens, including trait names, abbreviations, units, and symbology. Trait differences among parental species and hybrids based on results from our study indicating whether hybrids showed intermediate values or resembled one parent. N.S = no significant differences across groups.

| Trait | Units | Symbology | *P. balsamifera* | Hybrids | *P. trichocarpa* |
| --- | --- | --- | --- | --- | --- |
| Adaxial guard cell length (G<sub>U</sub>) | μm |  | N.S | N.S | N.S |
| Abaxial guard cell length (G<sub>L</sub>) | μm |  | Greater | *balsamifera*-like | Smaller |
| Adaxial pore length (P<sub>U</sub>) | μm |  | N.S | N.S | N.S |
| Abaxial pore length (P<sub>L</sub>) | μm |  | Greater | *balsamifera*-like | Smaller |
| Adaxial stomatal density (D<sub>U</sub>) | mm<sup>2</sup> |  | None or less dense | *trichocarpa*-like | Denser |
| Abaxial stomatal density (D<sub>L</sub>) | mm<sup>2</sup> |  | Less dense | *balsamifera*-like | Denser |
| Total stomatal density (D<sub>T</sub>) | mm<sup>2</sup> |  | Less dense | *balsamifera*-like | Denser |
| Stomata density ratio (SR) | None |  | More hypostomatous | *trichocarpa*-like | Higher degree of amphistomy |
| Adaxial stomatal occurrence (O<sub>U</sub>) | None |  | Lower | *trichocarpa*-like | Higher |
| Stomatal conductance (g<sub>sw</sub>) | mol m<sup>-2</sup> s<sup>-1</sup> |  | Higher | Intermediate values | Lower |
| Intrinsic water-use efficiency (δ13C) | ‰ | High/Low WUE | Lower | *balsamifera*-like | Higher |

*P. balsamifera* had significantly lower total stomatal density (D_T_) than *P. trichocarpa* (*p* < 0.05; Fig. 2A). Hybrid genotypes did not differ significantly from *P. balsamifera* (*p* > 0.05), but their D_T_ was significantly lower than *P. trichocarpa* (*p* < 0.05) and their range of phenotypic values exceeded both parental species. Abaxial stomatal density (D_L_) did not differ significantly between the parental species (*p* < 0.05), although *P. trichocarpa* tended to exhibit higher D_L_ than *P. balsamifera* (Fig. S3). Hybrid genotypes had significantly lower D_L_ than *P. trichocarpa* (*p* < 0.05) but did not differ from *P. balsamifera* (*p* > 0.05). Compared to both parental species, the range of hybrid D_L_ values was greater. *P. trichocarpa* exhibited significantly greater adaxial stomatal occurrence (O_U_) and density (D_U_) than *P. balsamifera* (*p* < 0.05, Fig. 2A). Hybrid genotypes did not differ significantly from *P. trichocarpa*, with greater O_U_ and a comparable range of traits values. *P. trichocarpa* exhibited a more even distribution of stomata across leaf surfaces, reflected in a significantly higher stomatal ratio (SR) compared to *P. balsamifera*, which had most stomata restricted to the lower surface (*p* < 0.05, Fig. 2A). Hybrids resembled *P. trichocarpa*, with a higher SR (*p* > 0.05) and similar range of values. *P. trichocarpa* showed significantly lower stomatal conductance (g_sw_) and higher water-use efficiency (δ¹³C) than *P. balsamifera* (*p* < 0.05; Fig. 2A). In hybrids, g_sw_ was intermediate between parents but exhibited a broader range of trait values. In contrast, δ¹³C values for hybrids more closely resembled *P. balsamifera* (*p* > 0.05) and the range did not exceed that observed in both parentals.

### Transect-specific climatic gradients influenced stomatal traits despite high interspecific gene flow

Geographic cline analyses suggested limited barriers to gene flow for stomatal traits (Fig. S4-S5), yet these traits were consistently associated with climatic gradients of origin, particularly precipitation (PC2) across the six latitudinal contact zones (Fig. 2B, Table S3). Opposing trait– climate relationships were observed between northern and southern contact zones for D_U_, O_U_, and SR. Drier environmental origins (PC2) were associated with significant decreases in D_U_, O_U_, and SR in the northern contact zones of Alaska, Cassiar, and Chilcotin, but significantly increased along PC2 in the southern contact zones of Jasper, Crowsnest, and Wyoming (Fig. 2B, Table S3, *p* < 0.05). Genotypes from warmer environments (PC1) were associated with increased D_U_ and lower O_U_ across the Jasper and Chilcotin contact zones, but reduced density and occurrence in the Wyoming contact zone (*p* < 0.05), while no significant associations were observed in the remaining contact zones (Fig. S6). Neither precipitation nor temperature gradients of origin were significantly associated with D_L_ or D_T_ across most contact zones (*p* > 0.05, Fig. S6), except for Alaska and Jasper, where warmer and drier origins were associated with increased D_T_ (*p* < 0.05). Together, these results suggest that transect-specific selection associated with climatic gradients have influenced density and distribution of stomata across the species’ contact zones.

Genotypes from drier origins (PC2) were associated with reduced G_L_ and P_L_ across most contact zones (*p* < 0.05), except for Alaska and Wyoming, where similar, but not significant, associations were observed (*p* > 0.05, Fig. 2B). In contrast to the abaxial surface, G_U_ and P_U_ did not vary significantly with precipitation gradients from site of origin (*p* > 0.05, Fig. S6) suggesting that stomatal size on the upper surface is influenced less by precipitation. Temperature gradients (PC1) were not significantly associated with guard cell length or pore length on either the adaxial or abaxial surface (*p* > 0.05, Fig. S6). These patterns suggest that variation in stomatal size across contact zones is more strongly influenced by precipitation than by temperature.

In contrast to stomatal density and size traits, g_sw_ was positively associated with temperature (PC1) and precipitation (PC2) gradients of origin across multiple contact zones (Fig. 2B, Table S3). Genotypes from cooler environments exhibited significantly greater g_sw_ (*p* < 0.05) with exception of Alaska and Cassiar. However, in Alaska, genotypes from drier environments had higher g_sw_ (*p* < 0.05). Although no significant associations were detected between climatic gradients and water-use efficiency (δ¹³C) in most contact zones, warmer genotype origins tended to be associated with reduced δ¹³C (Fig. S6). Drier genotype origins were negatively associated with δ¹³C in Cassiar and Wyoming (*p* < 0.05), but not in other contact zones (Fig. S6, Table S3), suggesting a trend toward lower water-use efficiency in drier environments for these contact zones.

### Admixture mapping reveals common and trait-specific candidate genes

Admixture mapping identified multiple ancestry blocks significantly associated with stomatal trait variation in hybrid genotypes (Table S4), revealing both trait-specific and shared regions of genomic association. The strongest associations were found for D_U_, SR, and O_U_, which were all associated with the same genomic region on chromosome 15 spanning approximately 66.5 kb. Multiple loci within this region exceeded the admixture burden-corrected significance threshold (Fig. 3A), suggesting a shared genetic basis (e.g., pleiotropy) for these traits. The region contained several candidate genes (Fig. 3B), including *TWIST* (a homologue of *SPEECHLESS* in *Arabidopsis*, a regulator of stomatal development, Peterson *et al*., 2010; Song *et al*., 2023), *TRY* (controls adaxial leaf surface trichome patterning, Matías-Hernández *et al*., 2016), *AGP22* (involved in cell wall structure via arabinogalactan peptides, Seifert & Roberts, 2007), *SF-RPO* (a plastid RNA polymerase sigma factor involved in chloroplast gene expression), *DFA* (*dihydrofolate synthetase*, involved in folate and nucleotide biosynthesis), and *MTM1* (a mitochondrial carrier domain protein involved in metal ion homeostasis, Su *et al*., 2007). In addition, a region on chromosome 12 associated with both D_U_ and SR contained *EPR1*, an early phytochrome-responsive transcription factor (Kuno, 2003, Fig. 3B). A third region on chromosome 9, containing *NF-YC10*, a transcription factor that regulates ABA signaling and contributes to drought tolerance (Swain *et al*., 2017), was associated with SR (Fig. 3B). Although no other traits passed the genome-wide significance threshold, we discuss suggestive associations for several additional traits in the Supplemental Material.

**Figure 3.**
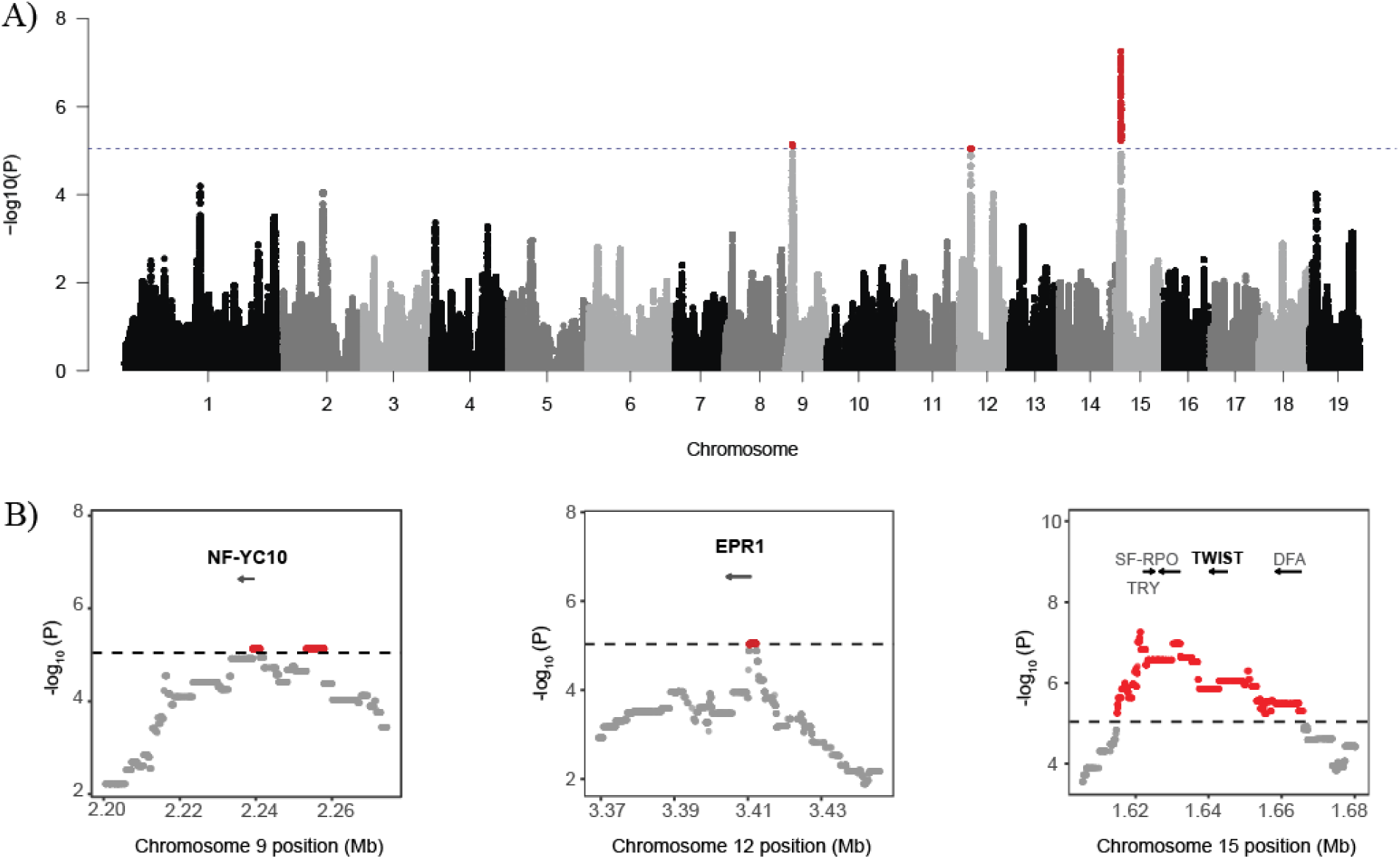
A) Manhattan plot showing results from GWAS by admixture mapping for stomatal ratio. The black dashed line indicates the genome-wide significance threshold (–log₁₀ (P) ≈ 4.73), corrected for admixture burden (0.05/2719). B) Zoomed-in views of significant associations with stomatal ratio on chromosomes 9, 12, and 15. Significant SNPs are highlighted in red. Candidate genes and their orientations are indicated by arrows.

### Transect-specific geographic clines suggests barriers to gene flow at candidate loci for stomatal traits

Geographic cline analyses revealed variation in cline centers and widths for candidate genes across the six *Populus* contact zones (Fig. 4, Table S5). In the Alaska contact zone, all stomatal candidate genes exhibited steep clines that mirrored genome-wide ancestry (*w*= 0.04, *c* = 0.64). Cline centers for these candidate genes were clustered towards the geographic distribution of *P. balsamifera* (*c* ≈ 0.65), and cline widths were narrow across candidate genes (*w* < 0.03). These patterns suggest that overall genetic exchange is restricted in this region. In contrast, geographic clines in Jasper were broader (*w* = 0.23–0.58), with centers ranging from *c* = 0.23 to 0.35 for both candidate genes and the genome-wide mean indicative of minimal barriers to gene flow.

**Figure 4.**
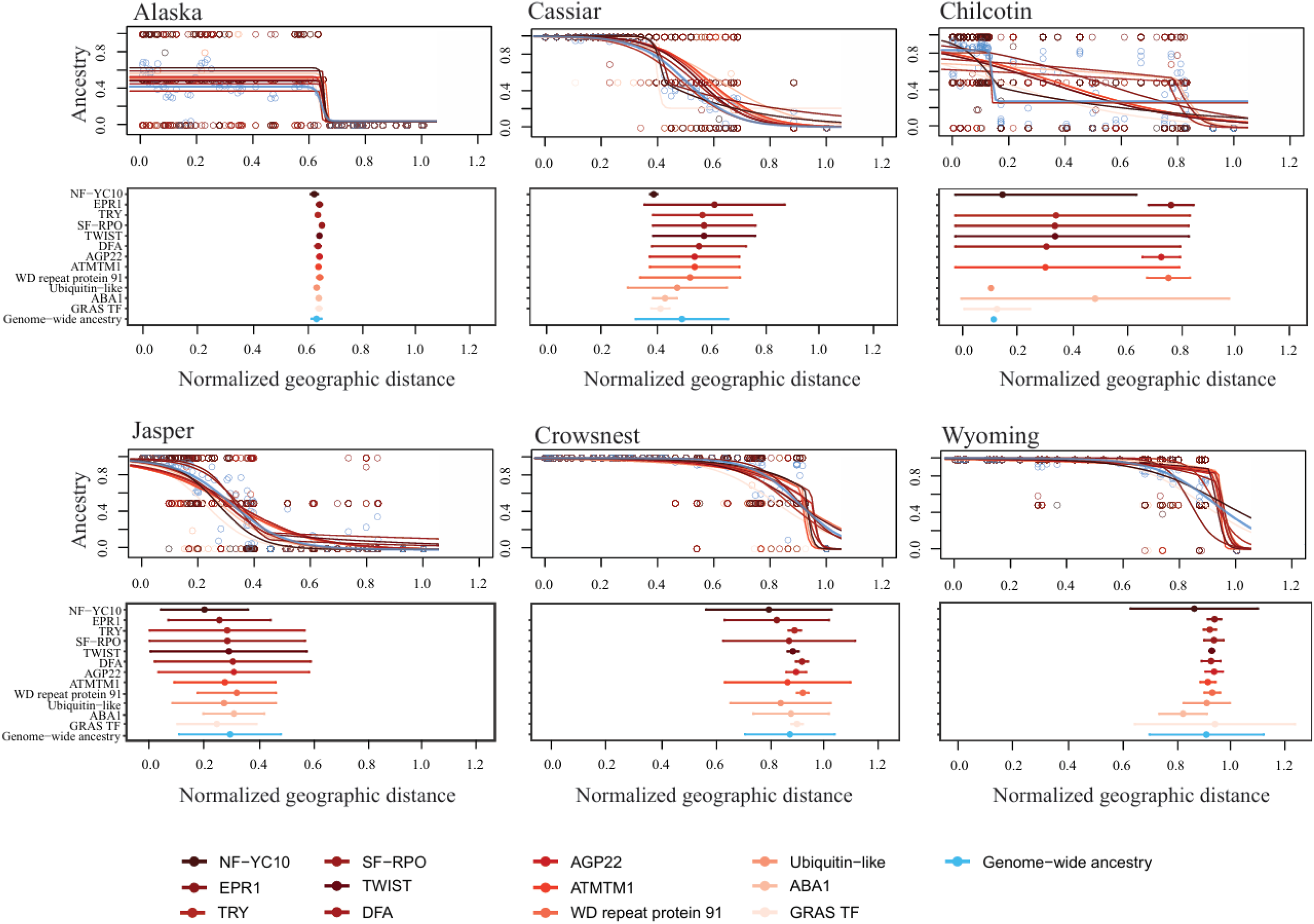
Geographic clines for candidate genes associated with stomatal trait variation and genome-wide ancestry across six contact zones in the *Populus balsamifera × P. trichocarpa* hybrid zone. Best-fit cline models estimated using HZAR illustrate the relationship between scaled geographic distance and ancestry (proportion of *P. trichocarpa* ancestry). Each colored line and dot represent a candidate gene, with cline shape determined by the model that best fits the observed ancestry frequency transition across the landscape. Lower panel provides a summary of cline parameters for each candidate gene. Points represent the estimated cline center (geographic midpoint of ancestry frequency change), and horizontal bars denote cline width (the spatial scale over which the transition occurs). Variation in cline shape, center, and width highlight differences in patterns of introgression and potential barriers to genetic exchange across transects.

In the Cassiar, Chilcotin, Crowsnest, and Wyoming contact zones, several candidate genes exhibited steeper clines when compared to the genome-wide background, indicating that potential barriers to gene flow have evolved at regions of the genome critical to species-specific adaptive trait variation (Fig. 4, Table S5). In Cassiar, the genome-wide ancestry cline was relatively broad (*w* = 0.34) and centered at *c* = 0.51. In contrast, several candidate genes associated with adaxial stomatal density, conductance, and guard cell size exhibited steeper and displaced clines (center differs from genome-wide cline), consistent with restricted introgression. For instances, *NF-YC10, ABA1,* and *GRAS transcription factor* exhibited steep clines (*w* = 0.03– 0.09) with centers (*c* = 0.40–0.44) consistently displaced toward the *P. trichocarpa* range, suggesting asymmetric introgression. In Chilcotin, the genome-wide ancestry cline was extremely narrow (*w* = 0.02), but clines at candidate genes varied widely, reflecting candidate gene-specific patterns of introgression. Several genes associated with adaxial stomatal density, *TRY, TWIST,* and *DFA,* exhibited broad clines (*w* = 1.00), suggesting extensive genetic exchange with centers displaced toward the *P. trichocarpa* range (*c* = 0.38, 0.37, and 0.34, respectively). In contrast, *AGP22* (adaxial stomatal density) and *WD repeat protein 91* (total stomatal density) showed narrower clines (*w* = 0.14 and 0.16) with centers at *c* = 0.77 and 0.80, respectively, consistent with localized barriers to introgression and displacement toward the *P. balsamifera* range. In Crowsnest, candidate genes associated with adaxial stomatal density including *TRY* (*w* = 0.05), *TWIST* (*w* = 0.04), *DFA* (*w* = 0.05), and *AGP22* (*w* = 0.08) exhibited narrower clines than the genome-wide average (*w* = 0.33). These genes had cline centers similar to the genome-wide center (*c* = 0.90), ranging from 0.91 to 0.95. Likewise, *WD repeat protein 91* (total stomatal density, *w* = 0.04, *c* = 0.95) and a *GRAS transcription factor* (upper guard cell size, *w* = 0.04, *c* = 0.93) showed similarly steep clines with centers also located within the *P. balsamifera* range, consistent with restricted introgression at genes influencing stomatal traits. In Wyoming, the genome-wide ancestry cline was relatively broad (*w* = 0.42), but most candidate genes exhibited much narrower transitions, with cline widths ranging from 0.02 to 0.18. With the exception of the *GRAS transcription factor* (*w* = 0.59) and *NF-YC10* (*w* = 0.47), which displayed cline widths comparable to or broader than the genome-wide pattern, most genes showed evidence of stronger barriers to gene flow. Cline centers were largely consistent across candidate genes, ranging from 0.87 to 0.95, and closely aligned with the genome-wide center (*c* = 0.92), with the exception of *ABA1*, which exhibited a slightly lower center (*c* = 0.84). Collectively, these results suggest that divergence is maintained at specific candidate genes involved in stomatal structure and function, but the extent and distribution of these barriers vary across contact zones, potentially due to differences in transect-specific selective pressures.

### Climatic associations suggest extrinsic selection shape local ancestry at genes associated with stomatal traits

Precipitation gradients associated with climate of origin had the strongest influence on local ancestry at stomatal trait candidate genes across contact zones (Fig. 5, Table S6). Precipitation gradients (PC2) were associated with reduced *P. trichocarpa* local ancestry (*p* < 0.01) at most candidate genes across Alaska, Cassiar, Chilcotin, Jasper, and Crowsnest contact zones (Fig. 5-S11). The probability of *P. trichocarpa* ancestry declined in genotypes originating from drier environments (*slopes*: –0.25 to –4.2), highlighting strong climate-mediated selection favoring *P. balsamifera* alleles where water availability was reduced. In Cassiar, genes associated with adaxial stomatal density and conductance, such as *ABA1*, *SF-RPO*, *TRY*, and *TWIST*, exhibited the steepest declines in *P. trichocarpa* ancestry along precipitation gradients (*slopes* = -2.8 to -4.23).

**Figure 5.**
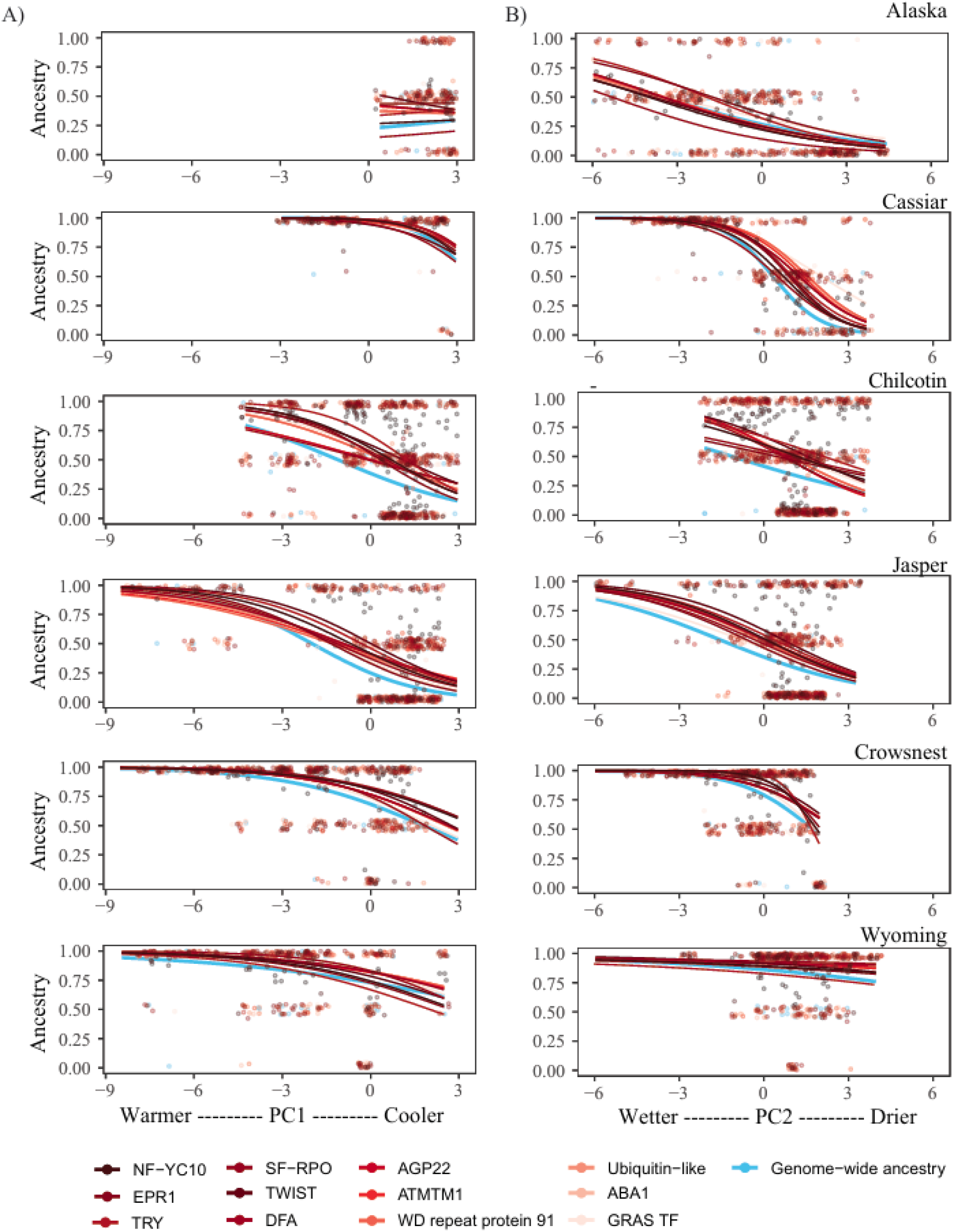
Relationship between climate and nuclear ancestry at stomatal trait candidate genes across six *Populus* contact zones. (A) Ancestry as a function of PC1, representing a gradient from warmer (low values) to cooler (high values) environments. (B) Ancestry as a function of PC2, representing a gradient from wetter (low values) to drier (high values) environments. Each point represents an individual genotype, colored by gene. Lines represent fitted values from generalized linear models (GLMs), fitted separately for each gene within each contact zone, with ancestry coded as 0 (*P. balsamifera* homozygotes), 0.5 (heterozygotes), or 1 (*P. trichocarpa* homozygotes).

Temperature gradients (PC1) based on genotype origin were significantly negatively associated with local ancestry for candidate genes across several contact zones, although these effects were generally weaker and less consistent than those driven by precipitation (PC2, Fig. 5-S11). In the Chilcotin, Jasper and Crowsnest contact zones, colder environmental origins were consistently associated with reduced presence of *P. trichocarpa* alleles at candidate genes, with slopes ranging from –0.27 to –1.29. In Crowsnest, genes associated with adaxial stomatal density and conductance, such as *ABA1*, *SF-RPO*, *TRY*, and *TWIST*, exhibited the steepest declines in *P. trichocarpa* ancestry along temperature gradients (*slopes* = -0.66 to -1.29). In Alaska and Cassiar, temperature gradients had limited influence on local ancestry, with only a few genes (*NF-YC10*, *AGP22*, *ATMTM1*, *ABA1*) showing significant but moderate declines in *P. trichocarpa* ancestry in cooler environments (slopes: –0.33 to –1.06).

Local ancestry at stomatal trait candidate genes was influenced by the interaction between temperature and precipitation gradients from the climate of origin in the Cassiar and Chilcotin contact zone (Fig. S11, Table S6). In Cassiar, genes involved in stomatal conductance and patterning such as *ABA1*, *AGP22*, and *TWIST* exhibited significant negative PC1 × PC2 interactions (*p* < 0.05), indicating selection for *P. balsamifera* ancestry under cold and dry conditions. In Chilcotin, similar interactions at multiple genes, including *NF-YC10*, *EPR1*, and *GRAS TF*, were found. Interestingly, when individual candidate genes were compared with genome-wide ancestry, consistent declines in *P. trichocarpa* allele frequencies were observed under colder and drier conditions across the six contact zones (Fig. 5).

## Discussion

As climate change accelerates, understanding how gene flow and selection interact to shape the evolution of adaptive traits will be critical to guiding species management. In this study, we investigated how interspecific gene flow and climate-mediated selection influence the evolution of stomatal traits that are crucial to plant growth, survival, and water-use efficiency, across six natural contact zones between *Populus trichocarpa* and *P. balsamifera*. Hybrid genotypes exhibited intermediate or parent-like phenotypes for key stomatal traits, but there was also substantial genetic variation mediated via hybridization. At a trait scale, geographic cline analyses indicated high genetic exchange, however, analysis of candidate genes underlying stomatal traits showed steep clines aligned with climatic gradients. Such gene-specific patterns indicate that extrinsic selection plays an important role influencing the movement of specific alleles across environmental gradients even in the presence of species barriers associated with adaptive trait differences. Together, our results provide new insight into how extrinsic selection impacts the movement of genetic variants into novel genomic and environmental backgrounds that may be critical to adaptive trait evolution.

### Species-specific differences in stomatal traits influence hybrids

Stomatal trait differences between *P. balsamifera* and *P. trichocarpa* reflect evolutionary responses to their home environments. In this study, traits associated with water conservation, such as reduced stomatal conductance, a high density of small stomata, and increased water-use efficiency were observed in *P. trichocarpa* compared to *P. balsamifera*. This contrasts with earlier findings that associated *P. balsamifera* with more water conservation strategies (Pointeau & Guy, 2014). One possibility is that *P. trichocarpa* may have experienced lower water potentials in the common gardens, leading to stomatal closure to avoid cavitation, consistent with a more drought-sensitive strategy. In contrast, *P. balsamifera* may have maintained turgor and kept stomata open under mild stress, reflecting greater drought tolerance (Groover *et al*., 2025). Differences may also arise from broader geographic sampling in our study, which captured more intraspecific variation than previous studies (Pointeau & Guy, 2014). However, patterns of amphistomy were consistent with previous findings in which *P. balsamifera* generally exhibited none or fewer stomata on the adaxial leaf surface, while *P. trichocarpa* showed greater variation, with more genotypes displaying adaxial stomata (McKown *et al*., 2014, 2019; Fetter *et al*., 2021).

Stomatal traits for hybrid genotypes were on average, intermediate or more similar to one of the parental species but showed greater ranges of phenotypic values suggesting that hybridization has generated a wide spectrum of trait variation upon which selection may act. Stomatal conductance in hybrids on average was intermediate to *P. balsamifera* and *P. trichocarpa*, consistent with additive genetic contributions from both parental species (Orians, 2000). Such additive trait expression may offer functional benefits in transitional environments where hybrids encounter conditions not occupied by either parental species (Janes & Hamilton, 2017). In our study, hybrid genotypes largely resembled *P. trichocarpa* in adaxial stomatal density, occurrence, and stomatal ratio with the presence of stomata on both leaf surfaces. Previous studies have reported adaxial stomata in *P. balsamifera* admixed genotypes only in regions of hybridization with *P. trichocarpa*, *P. deltoides*, or *P. angustifolia* (McKown et al. 2014, 2019; Pearce et al., 2005), suggesting that amphistomaty arises through introgression. In hybrids, amphistomaty may provide an advantage, enhancing photosynthetic capacity across transitional environments (Muir, 2019). In contrast, in *P. deltoides × P. angustifolia* hybrids, amphistomatous traits from *P. deltoides* were not retained, and both *P. angustifolia* and hybrids remained largely hypostomatous (Zanewich et al., 2018). These contrasting outcomes suggest that the expression of amphistomaty in hybrids may be shaped by differences in parental trait dominance or directional selection (Pfeilsticker et al., 2022, 2023).

### Climate shapes genetic variation underlying stomatal traits despite high interspecific gene flow

Despite geographic clines indicating limited barriers to gene flow at a trait level across the six latitudinal contact zones, trait variation was strongly associated with climatic gradients, particularly precipitation. Adaxial stomatal density, occurrence, and stomatal ratio were strongly associated with precipitation gradients although the direction of the relationship varied across several contact zones. In the northern contact zones, wetter origins were associated with higher adaxial stomatal density with more genotypes with amphistomatous leaves, while drier origins showed reduced adaxial stomata and more genotypes with hypostomatous leaves. This suggests that, in the north, selection may favor amphistomaty under wetter conditions to support rapid growth during short growing seasons, but is selected against in drier climates to limit water loss (McKown *et al*., 2019). Southern contact zones showed the opposite pattern with lower stomatal ratio for genotypes sourced from wetter environments and higher from drier origins. In the south, selection may favor lower adaxial stomatal density to reduce pathogen entry, reflecting a trade-off between gas exchange and disease resistance which is typically higher in warmer, humid regions of the south compared to the north (McKown *et al*., 2014; Fetter *et al*., 2021).

Stomatal size traits, including guard cell and pore length, showed clear associations with precipitation across the hybrid zone. In most contact zones, genotypes from drier environments were linked to smaller abaxial guard cells and shorter pore lengths, supporting the hypothesis that reduced stomatal size reduces water loss under drought stress while allowing faster stomatal closure (Niemczyk *et al*., 2019; Volk *et al*., 2022; Franks & Beerling, 2009). Contrary to expectations that genotypes originating from cooler and drier climates would exhibit reduced stomatal conductance (Chen *et al*., 2024), we observed that these genotypes maintained higher stomatal conductance when evaluated in common gardens. Lower evaporative demand in cooler environments may reduce the cost of water loss, allowing higher stomatal conductance with minimal risk. In drier regions, genotypes may exploit brief water availability by maximizing carbon gain during short favorable periods (Buckley, 2019). However, observed patterns may also reflect environmental differences between the common gardens and the climates of origin (Oubida *et al*., 2015).

### Climate-driven selection limits gene flow at loci underlying stomatal traits

Although trait-scale geographic clines suggested minimal barriers to genetic exchange for stomatal traits, finer-scale cline analyses of candidate genes underlying stomatal traits revealed variable restrictions to gene flow, depending on genomic context and environmental conditions. Most contact zones exhibited evidence of gene-specific constraints to genetic exchange; however, the Alaska contact zone with coincident patterns of limited introgression for adaptive and neutral loci may reflect broad ecological constraints. In this contact zone, the cline for genome-wide ancestry was extremely narrow (*w* = 0.04) and all candidate genes showed similarly steep transitions (*w* < 0.03), suggesting barriers to gene flow reflect genome-wide divergence (Schield *et al*., 2024). While steep coincident clines can also emerge from strong genetic drift (Jofre & Rosenthal, 2021), the consistent decline in *P. trichocarpa* ancestry toward drier climates (Fig. 5) supports the idea that precipitation gradients are shaping local ancestry at candidate genes for stomatal traits. Similar studies in hybrid zones have shown that precipitation can act as a key selective force influencing patterns of genetic variation and introgression (Hamilton & Aitken, 2013; Menon *et al*., 2021).

In contrast to Alaska, the Cassiar, Chilcotin, Crowsnest, and Wyoming contact zones exhibited more porous species boundaries, where barriers to gene flow were locus-specific and strongly influenced by climate of origin. In Cassiar, clines at candidate genes associated with adaxial stomatal density and conductance including *ABA1*, *NF-YC10*, and *TRY* were markedly steeper (*w* < 0.09) than the genome-wide average (*w* = 0.34), consistent with divergent selection limiting introgression at adaptive loci underlying traits that show strong species-specific divergence (Barton & Gale, 1993; Suarez-Gonzalez *et al*., 2018c). Notably, this fine-scale restriction contrasts with the broader clines (*w* = 0.33 -0.52) seen in other candidate genes and genome-wide ancestry (*w* = 0.35) indicating that while much of the genome can move freely across species boundaries, regions underlying traits like stomatal conductance and adaxial stomatal density are subject to divergent selection. Consistent with climate-driven selection, we found strong ancestry–climate associations, including significant temperature × precipitation interactions, suggesting steep climatic gradients may impact recombination in Cassiar. Evidence from chloroplast haplotype distributions in this zone also suggests introgression is climate-mediated (Zavala-Paez *et al*., 2025). These results highlight the role of extrinsic selection in maintaining species differences where there might otherwise be weak barriers to reproduction (Abbott, 2017). Unlike other contact zones, Jasper showed little evidence of barriers to gene flow, either genome-wide or at candidate genes. Clines at both genome-wide loci (*w* = 0.58) and candidate genes (*w* = 0.23–0.58) were broad, indicating gene flow and a lack of reproductive isolation. Yet, consistent climate associations, especially with precipitation, suggest that selection is acting, but currently the strength is insufficient to overcome the homogenizing effect of gene flow.

Taken together, comparisons across the six latitudinal contact zones reveal that climate-mediated selection, particularly variation in precipitation, constrains introgression both across the genome and at functionally important candidate genes. However, the strength of this selection varies with local environmental conditions, resulting in spatial differences in allele movement. Despite widespread interspecific hybridization, our findings demonstrate that environmental selection can maintain divergence at key loci. These results offer insight into the mechanisms that both limit and facilitate the movement of adaptive alleles, ultimately shaping the evolution of adaptive traits in hybrid zones.

## Conclusion

This study reveals that stomatal trait evolution in *Populus* hybrid zones is shaped by the interplay of gene flow and climate-driven selection. Hybrids displayed a wide range of trait values, reflecting admixture between divergent parental species. Variation in stomatal traits was strongly associated with environmental gradients, particularly precipitation, suggesting that water availability plays a central role in influencing trait variation across this hybrid zone despite high levels of gene flow. However, although much of the genome exhibits high levels of interspecific gene flow at a trait-scale, several candidate genes associated with amphistomy and stomatal conductance exhibited restricted gene flow. Variation in the movement of genetic variants underlying adaptive traits was largely influenced by climate of origin shaping local ancestry distributions. These findings highlight that hybridization can generate novel genetic recombinants and facilitate genetic exchange in areas where barriers to reproduction are limited. But overall, the movement of adaptive genetic variation across species and environmental gradients will be strongly influenced by environmental selection.

## Acknowledgments

We thank Baxter Worthing for assistance with trait sampling in the Vermont common garden, and Sara Klopf and Tommy Phannareth for their help with trait sampling in the Virginia common garden. We are also grateful to Kyle Peer, Clay Sawyers, and Deborah Bird at the Virginia Tech Reynolds Homestead Forestry Research Station for their support with plant propagation. Additional thanks go to Nadia Garzione for her contributions to stomatal trait sampling and processing. Finally, we appreciate Alayna Mead, Kyra LoPiccolo, and Sammy Muraguri from the Hamilton Lab for their insightful questions and engaging discussions during lab meetings, which helped shape the direction of this work.

## Author contributions

Michelle Zavala-Paez led data collection, research design, data analysis and interpretation, and manuscript writing. Stephen Keller contributed to data collection, data analysis and interpretation, and manuscript writing. Jason Holliday and Matthew Fitzpatrick contributed to data collection and manuscript writing. Jill Hamilton contributed to data collection, research design, data analysis and interpretation, and manuscript writing.

## Data availability

The genomic data analyzed in this study were previously published by Bolte et al. (2024) and are available on NCBI under accession number PRJNA996882. All data and scripts used for the analyses will be deposited in Dryad upon acceptance and are provided in the supplementary materials for reviewers.

