## Supplementary material for "Climate and hybridization shape stomatal trait evolution in *Populus*": Supplementary_Material.docx

**Supplementary methods**

### **Geographic clines for stomata traits and associated loci**

Geographic clines for stomatal traits and associated loci were constructed to test the extent and direction of introgression of the genetic variation underlying stomatal traits. Geographic distances within each contact zone were calculated using the Haversine formula, based on the latitude and longitude coordinates of individual genotypes. For each genotype, geographic distance $D_{norm}$ was normalized using the following equation:

$$D_{norm}= \frac{D_{i}- D_{min}}{D_{max}+ D_{min}}$$

Where: $D_{i}$ is the geographic distance for genotype *i*, $D_{min}$ is the minimum geographic distance in the contact zone, and $D_{max}$ is the maximum geographic distance in the contact zone. To facilitate direct comparisons across contact zones, geographic distances were normalized to a scale from 0 to 1.

**Supplementary results**

**Putative genomic associations with stomatal trait variation**

Admixture mapping analysis showed suggestive associations (i.e., below the threshold but potentially biologically meaningful) for several stomatal traits (Fig. S7-S10). For adaxial guard cell length, a region on chromosome 7 included a *GRAS family transcription factor* (involved in gibberellin-mediated cell elongation). For abaxial guard cell length, a region on chromosome 6 contained Potri.006G059700, encoding a *ubiquitin-like protein*. For total stomatal density, a region on chromosome 15 included *Potri.015G026100*, a *WD repeat-containing protein 91* (involved in protein–protein interactions). Finally, for stomatal conductance, a region on chromosome 7 contained *ABA1*, a key enzyme in abscisic acid biosynthesis that regulates stomatal aperture.

**Supplementary Figures**

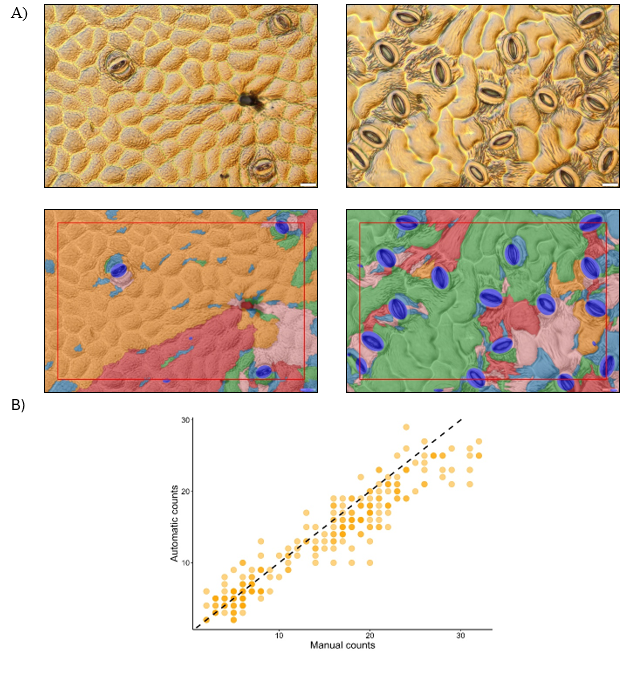

Figure S1. Comparison of manual and automated stomatal counts. A) Micrographs of epidermal impressions taken at 20 µm scale. LeafNet output highlights detected stomata with blue ovals. B) Spearman correlation (ρ = 0.95) between stomatal density estimates obtained manually and using LeafNet, with a 1:1 reference line shown in black.

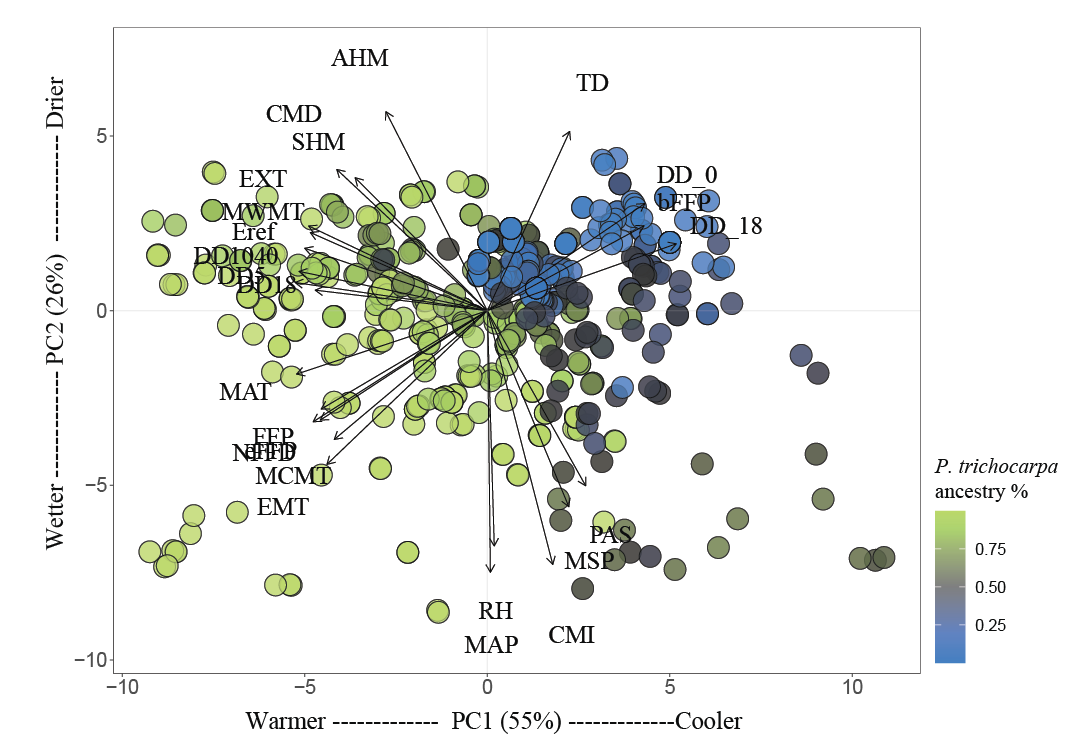

Figure S2. Principal component analysis (PCA) of 25 climate variables associated with genotype origin. PCA summarizes 25-year climate normals into two major axes. PC1 (55%) captured temperature-related gradients, with high loadings for mean annual temperature and degree-days above 5 °C and 18 °C. PC2 (26%) captured moisture-related gradients, with high loadings for mean annual precipitation, climate moisture index, and relative humidity. Points represent individual sampling sites colored *P. trichocarpa* ancestry variable. Climate vectors indicate variable loadings, with arrows pointing in the direction of increasing values.

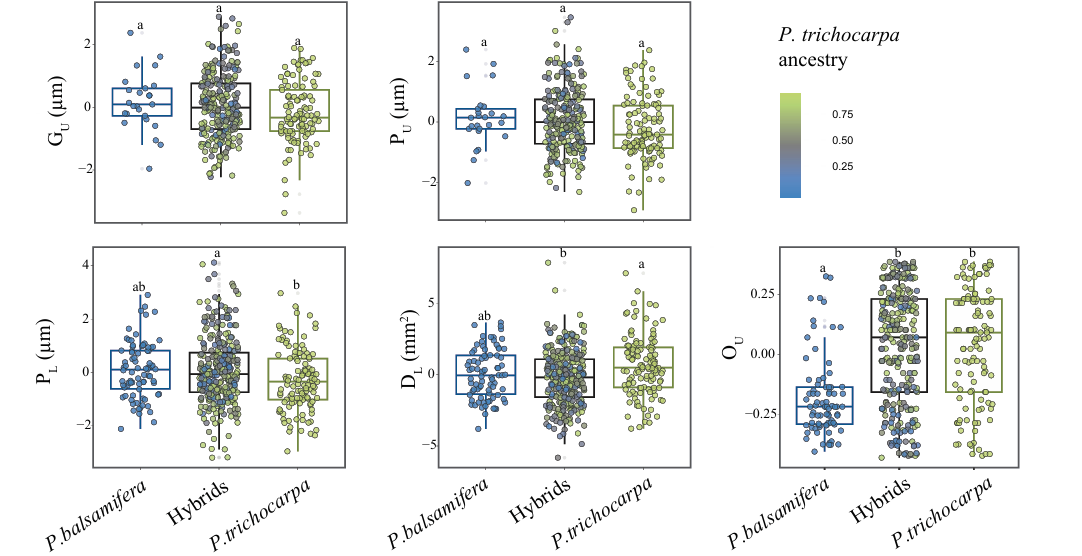

Figure S3. Boxplots show best linear unbiased predictors (BLUPs) for adaxial guard cell length (G_U_), adaxial pore length (P_U_), abaxial pore length (P_L_), abaxial stomatal density (D_L_), and abaxial stomatal occurrence (O_U_), for *P. balsamifera*, admixed, and *P. trichocarpa* genotypes. Each point represents a genotype, and colors indicate the proportion of *P. trichocarpa* genomic ancestry. Letters denote statistically significant differences among groups based on post-hoc comparisons (p < 0.05).

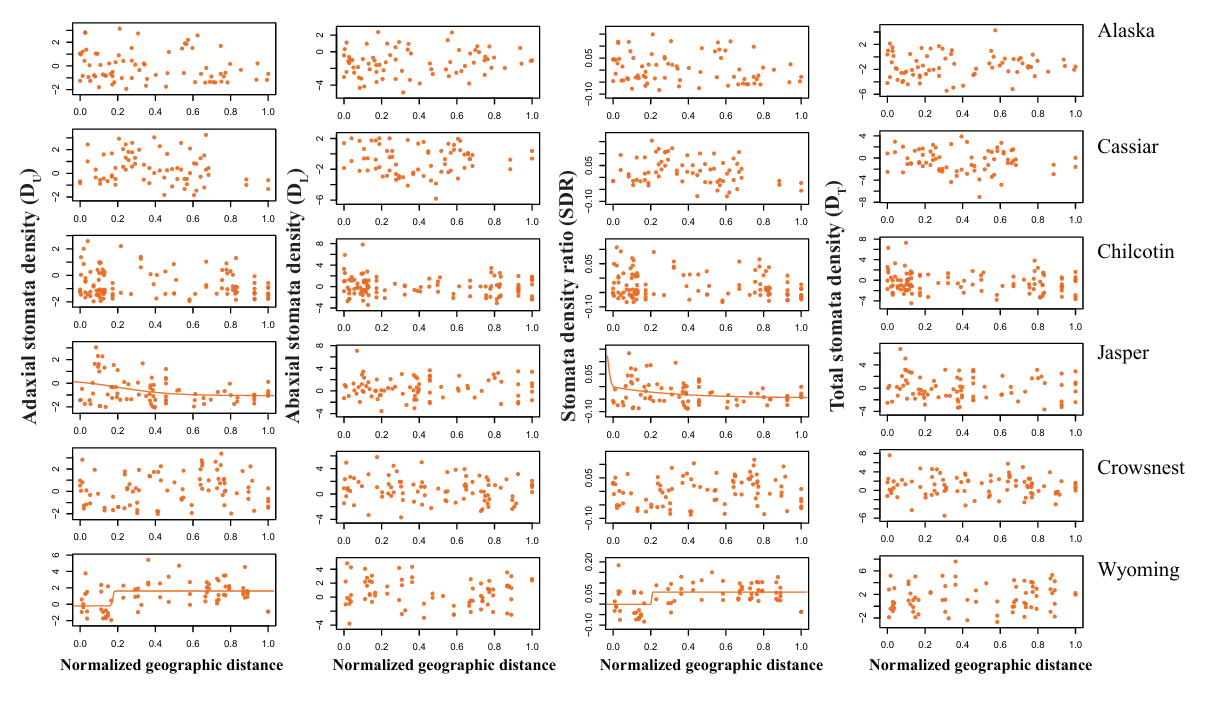

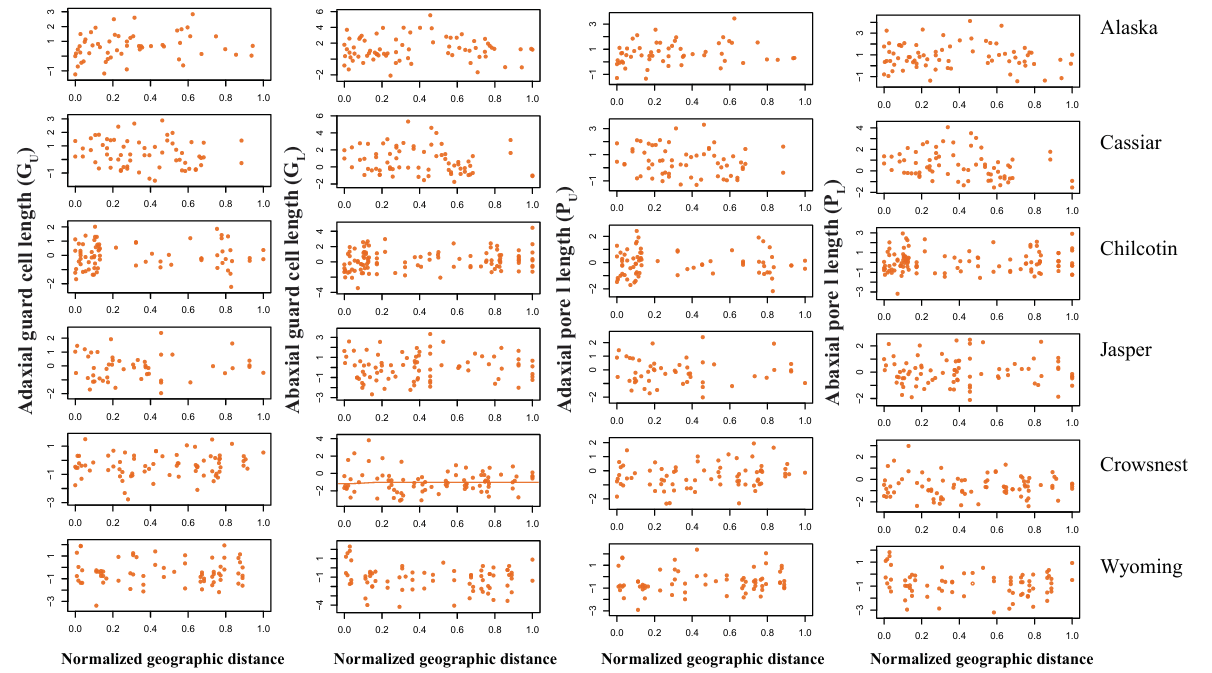

Figure S4. Relationship between normalized geographic distance and stomatal trait values. Points represent an individual genotype.

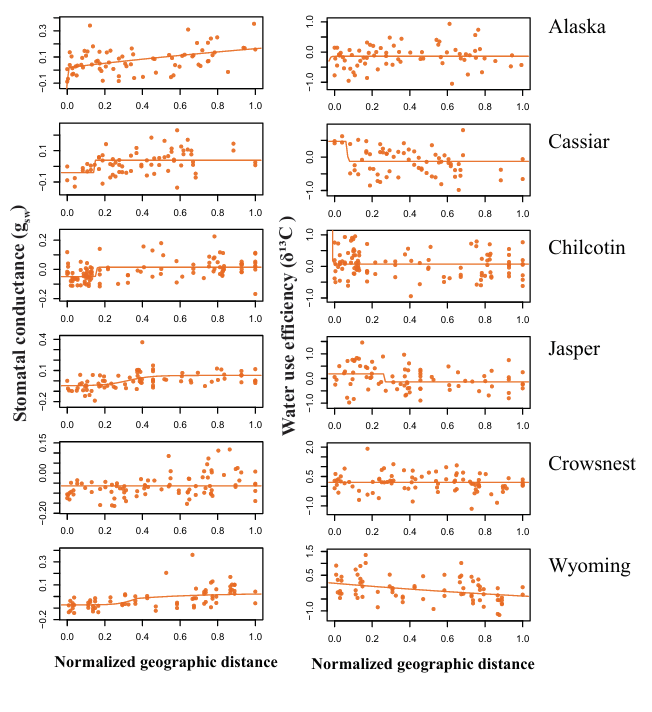

Figure S5. Relationship between normalized geographic distance and stomatal trait values. Points represent an individual genotype

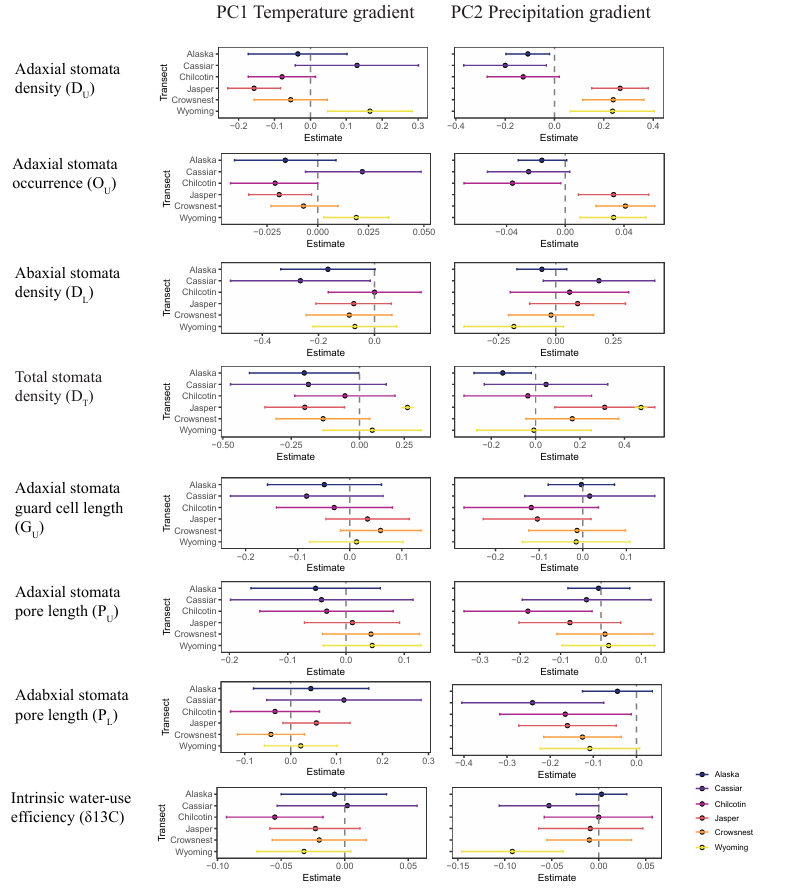

Figure S6. Relationship between genotype BLUPs values stomatal traits and the first two principal components of climate at the site of origin. PC1 reflects a gradient from warmer to cooler environments (positive values = cooler), while PC2 reflects a precipitation gradient (positive values = drier). Panels show trait-transects slopes (±95% CI) extracted from linear models fit separately within each transect. Positive slope values indicate trait increases along the climate axis (e.g., toward drier or warmer environments), while negative values reflect decreases.

Adaxial pore length (P_U_)

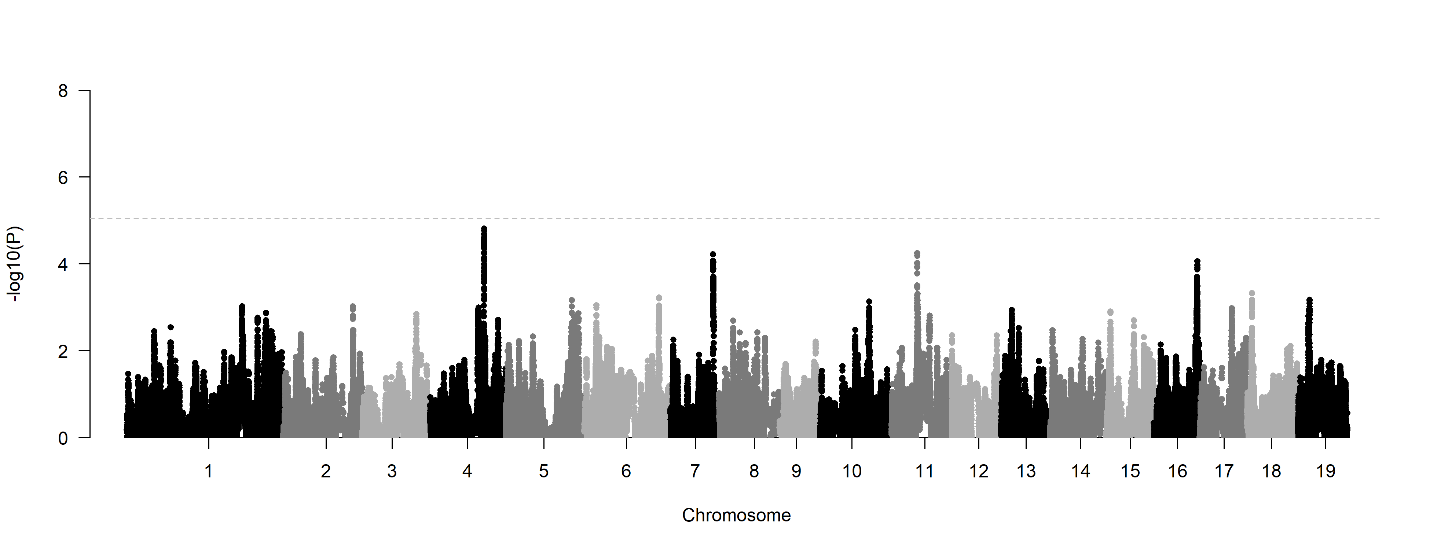

Abaxial pore length (P_L_)

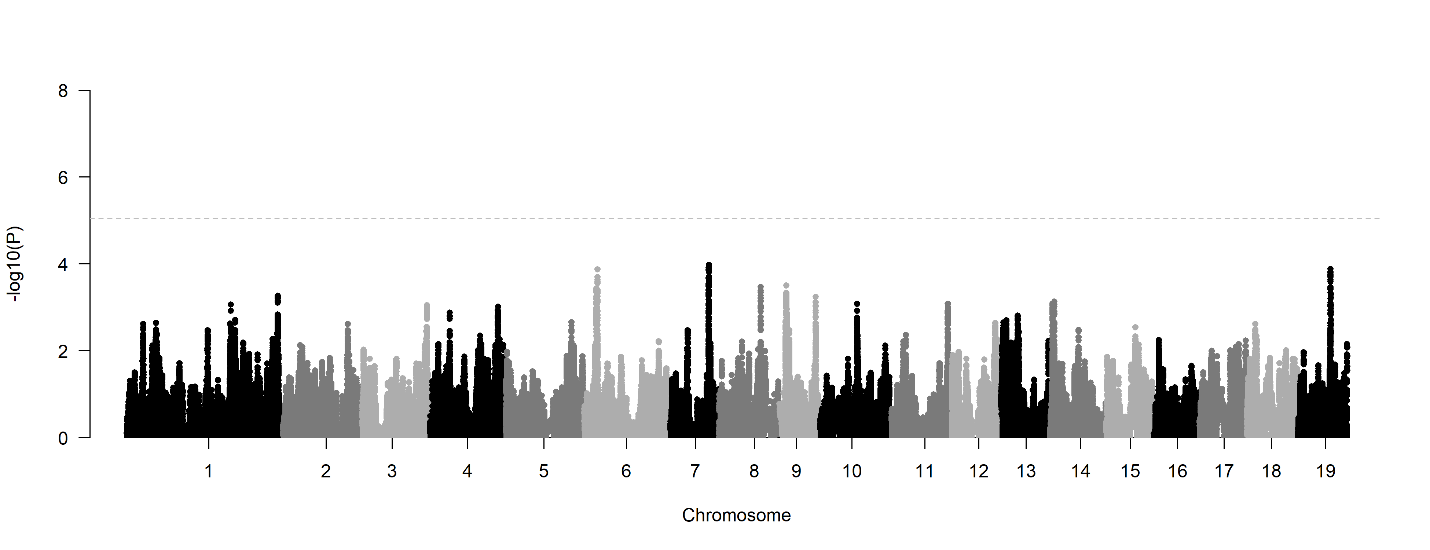

Figure S7. Manhattan plots showing results from GWAS by admixture mapping for adaxial and abaxial pore length. While no SNPs surpassed the genome-wide significance threshold (gray dashed line; –log₁₀(P) ≈ 4.73), several SNPs showed suggestive associations (i.e., below the threshold but potentially meaningful), including some located within genes or within 2 kb upstream

Abaxial stomatal density (D_L_)

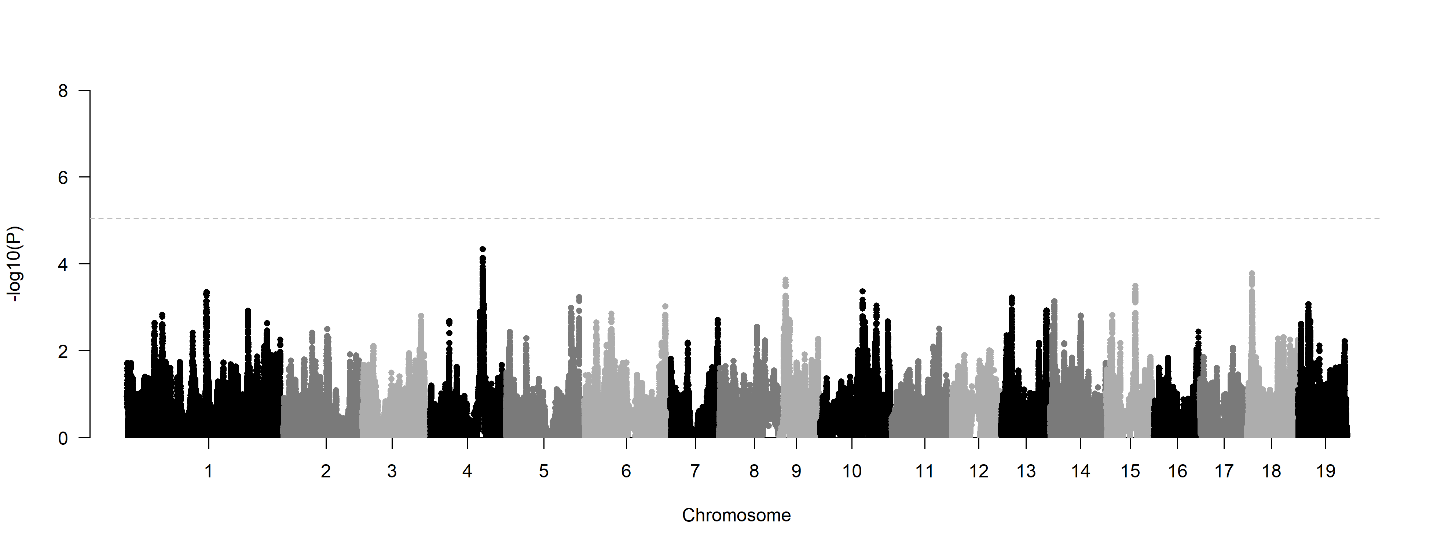

Intrinsic water-use efficiency (δ13C)

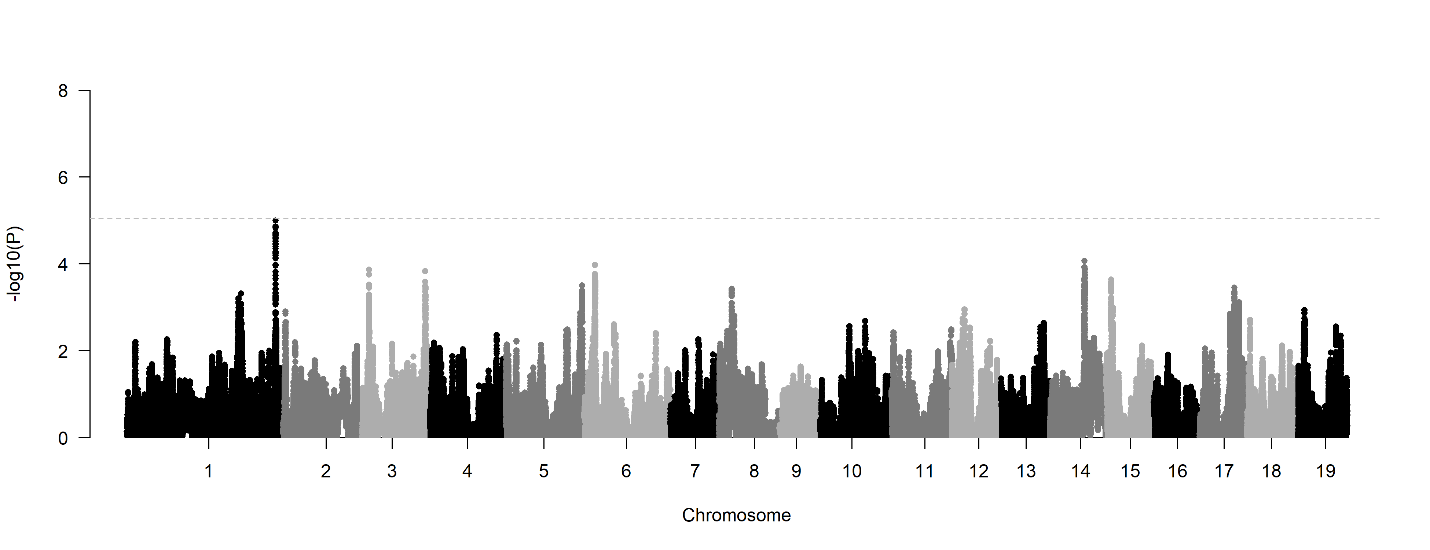

Figure S8. Manhattan plots showing results from GWAS by admixture mapping for abaxial stomatal density and intrinsic water-use efficiency. While no SNPs surpassed the genome-wide significance threshold (gray dashed line; –log₁₀(P) ≈ 4.73), several SNPs showed suggestive associations (i.e., below the threshold but potentially meaningful), including some located within genes or within 2 kb upstream.

Adaxial guard cell length (G_U_)

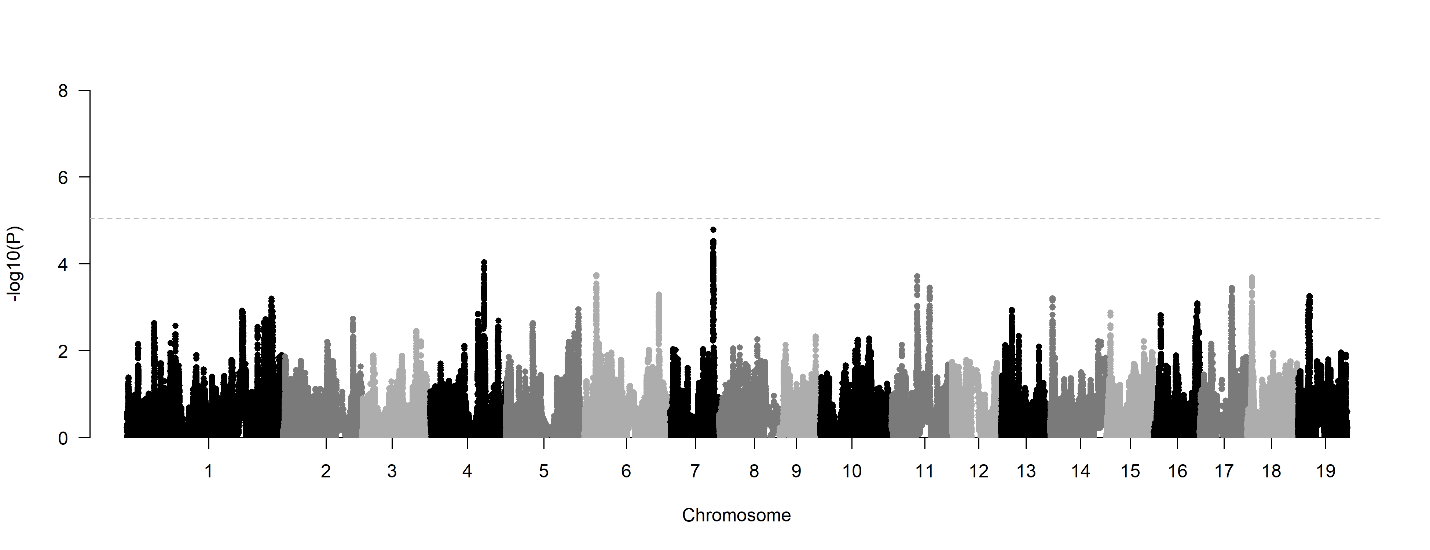

Abaxial guard cell length (G_L_)

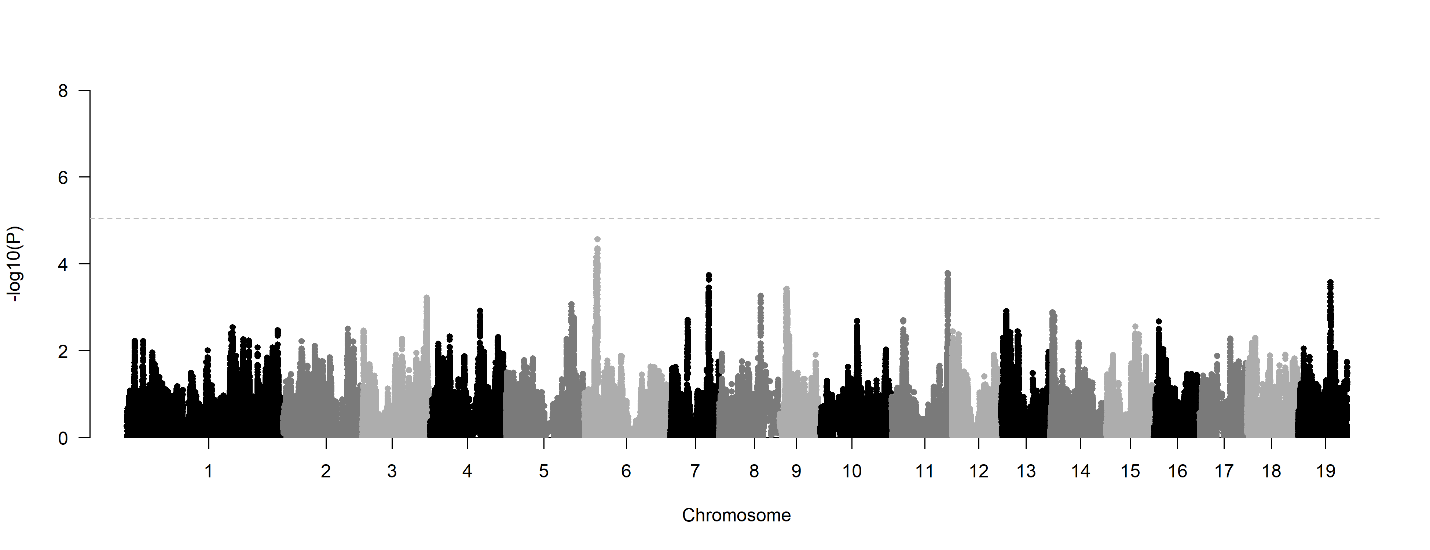

Figure S9. Manhattan plots showing results from GWAS by admixture mapping for adaxial and abaxial guard cell length. While no SNPs surpassed the genome-wide significance threshold (gray dashed line; –log₁₀(P) ≈ 4.73), several SNPs showed suggestive associations (i.e., below the threshold but potentially meaningful), including some located within genes or within 2 kb upstream.

Total stomatal density (D_T_)

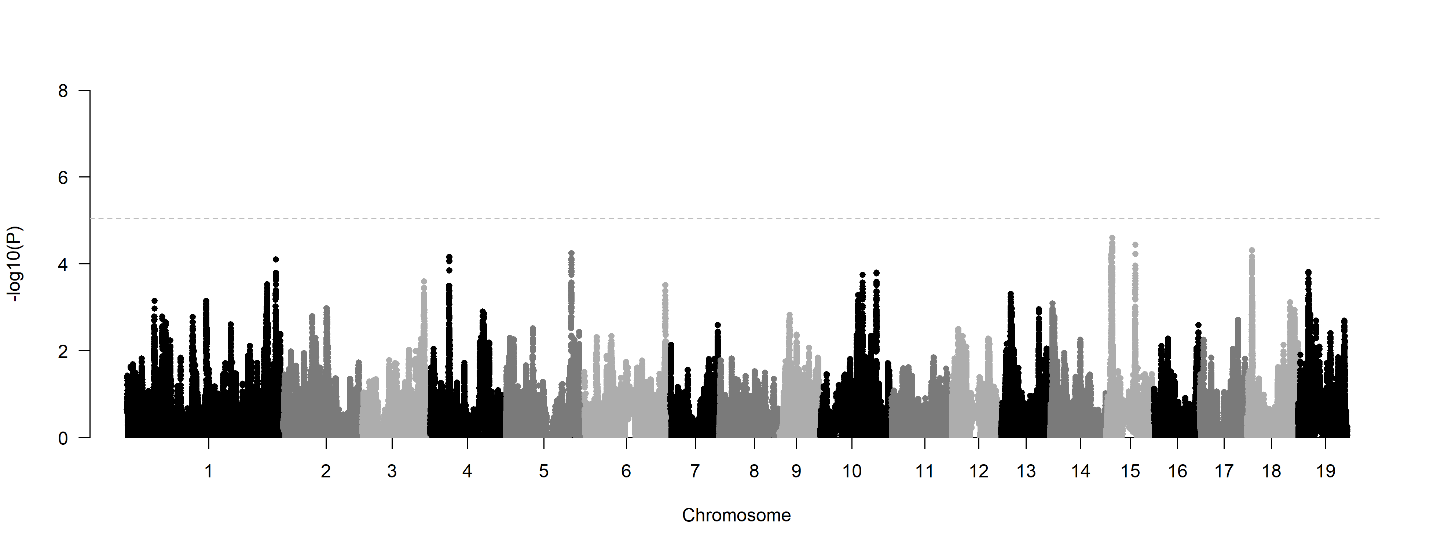

Stomatal conductance (g_sw_)

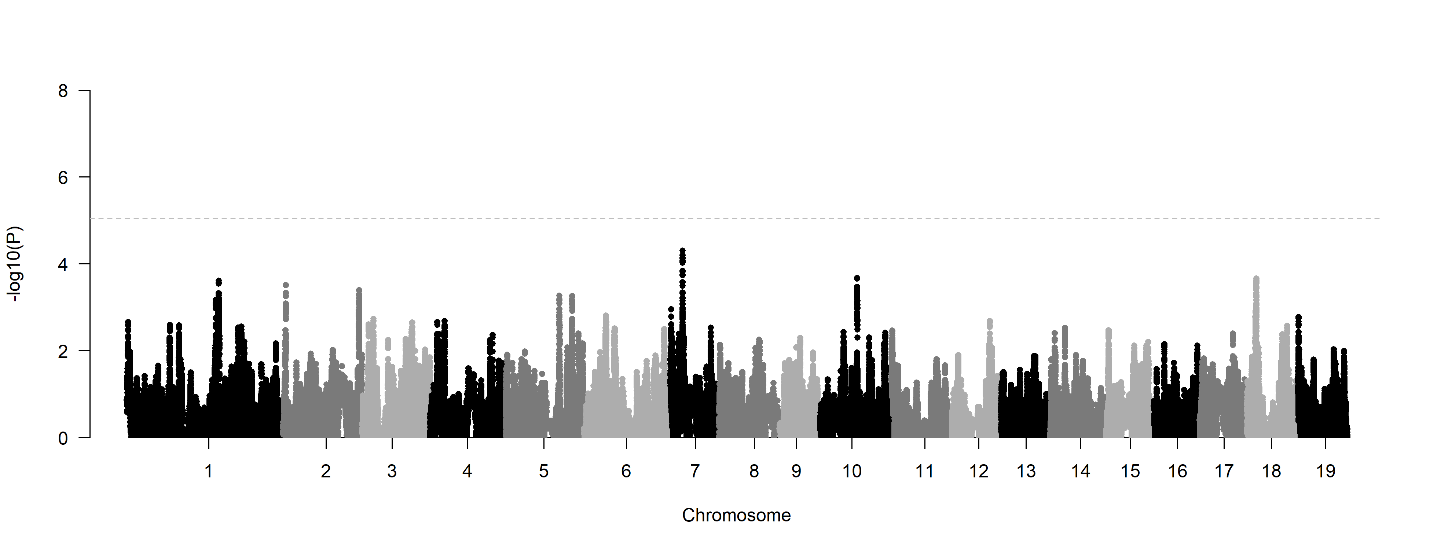

Figure S10. Manhattan plots showing results from GWAS by admixture mapping for total stomatal density and stomatal conductance. While no SNPs surpassed the genome-wide significance threshold (gray dashed line; –log₁₀(P) ≈ 4.73), several SNPs showed suggestive associations (i.e., below the threshold but potentially meaningful), including some located within genes or within 2 kb upstream.

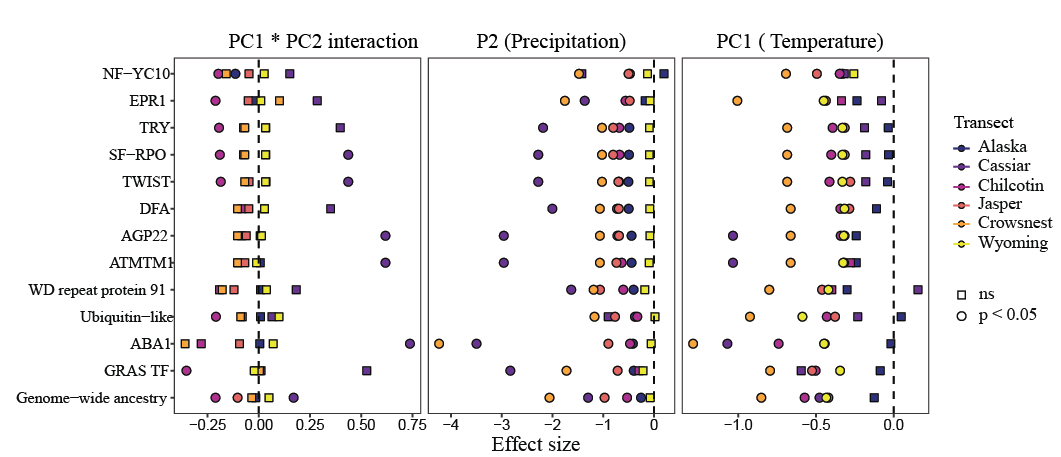

Figure S11. Effect sizes of climate variables on ancestry at stomatal trait candidate genes across six Populus hybrid zones. Effect sizes from quasibinomial GLMs are shown for the main effects of temperature (PC1), precipitation (PC2), and their interaction (PC1 × PC2). Each point represents the estimated effect for a gene in a specific transect, with fill color indicating transect. Circles indicate statistically significant associations (*p* < 0.05), and squares indicate non-significant associations.
